# Protein and Lipid Mass Concentration Measurement in Tissues by Stimulated Raman Scattering Microscopy

**DOI:** 10.1101/629543

**Authors:** Seungeun Oh, ChangHee Lee, Wenlong Yang, Ang Li, Avik Mukherjee, Markus Basan, Chongzhao Ran, Wei Yin, Clifford J. Tabin, Dan Fu, X. Sunney Xie, Marc W. Kirschner

## Abstract

Cell mass and its chemical composition are important aggregate cellular variables for physiological processes including growth control and tissue homeostasis. Despite their central importance, it has been difficult to quantitatively measure these quantities from single cells in intact tissue. Here, we introduce Normalized Raman Imaging (NoRI), a Stimulated Raman Scattering (SRS) microscopy method that provides the local concentrations of protein, lipid and water from live or fixed tissue samples with high spatial resolution. Using NoRI, we demonstrate that single cell protein, lipid and water concentrations are maintained in a tight range in cells under same physiological conditions and are altered in different physiological states such as cell cycle stages, attachment to substrates of different stiffness, or by entering senescence. In animal tissues, protein and lipid concentration varies with cell types, yet an unexpected cell-to-cell heterogeneity was found in cerebellar Purkinje cells. Protein and lipid concentration profile provides a new means to quantitatively compare disease-related pathology as demonstrated using models of Alzheimer’s disease. Our demonstration shows that NoRI is a broadly applicable tool for probing the biological regulation of protein mass, lipid mass and water in cellular and tissue growth, homeostasis, and disease.

---

All biological cells must tightly control their volume, mass, and molecular composition to ensure cellular fitness^1^ or functionality in a tissue context^2^. Cell mass and volume growth is closely coordinated with proliferation^3,4^ and tissue growth^5^, and the ratio of mass to volume, that is mass concentration, reflects macromolecular crowding of intracellular milieu^6^ and influences cellular fitness^1^. Despite their conceptual simplicity and importance, our understanding how cells regulate and coordinate these aggregate physiological properties has been hampered by the difficulty of accurately measuring them, especially in tissue contexts^7^. Technology to accurately determine single cell mass and volume has been both a driving force and a limitation for addressing these questions. For instance, several methods have been developed for single cell measurement of either cell volume^8–11^, cell mass^12–18^, or surrogate variables that correlate with cell size^4,19,20^ (Box 1). Despite the general assumption that cell mass and volume are proportional to each other, cell volume can dramatically deviate from cell mass^5^. As the importance of the macromolecular density in cytoplasm has been raised, a few technologies have been developed to quantitatively characterize the density of intracellular milieu^21,22^ and subcellular compartments, such as the phase separated compartments^23,24^. However, existing methods lack subcellular resolution, or are limited in their applicability in a tissue context. As a result, our current knowledge on the regulation of cell size and cytoplasmic density are drawn from bulk measurements, average behavior of cell populations, suspended or cultured cells *in vitro*, or from measurement of proxy variables that are bound to specific contexts. Moreover, current single cell mass measurements only provide total biomass, and cannot differentiate between protein and lipid mass.

To develop a method that overcomes the limitations of existing approaches, we turned to Stimulated Raman scattering (SRS) microscopy^25^, which is ideal for quantitative analysis of cellular materials in tissue. However, existing SRS approaches suffer from heterogeneous signal attenuation due to light scattering in thick samples, limiting their utility in accurate quantification of single cell mass *in situ*^26^. Here we report an alternative approach that enables accurate determination of single cell mass and cytoplasmic mass concentration in live or fixed tissue by computationally removing the effect of light scattering. The key step in NoRI is the conversion of the SRS images to absolute concentrations through normalization of the undesirable intensity variation caused by sample light scattering. Hence, we name this new method Normalized stimulated Raman scattering Imaging (NoRI). Use of the sum of chemical components including water is the key element in converting chemical compositions to absolute concentration in a way comparable to those demonstrated in other Raman modalities^27,28^.

## Box 1. Comparison of NoRI and existing methods for single cell biomass measurement

| Method | Measured parameters | Measurement characteristics |  |  |  |  | Sample types |  |  |  | References |
| --- | --- | --- | --- | --- | --- | --- | --- | --- | --- | --- | --- |
|  |  | Mass | Mass density | Absolute quantification* | Subcellular resolution | Chemical composition | Suspended cells | Adherent cell culture | Cells in tissue | Live sample |  |
| NoRI | Biomass composition and concentration | ✓ | ✓ | ✓ | ✓ | ✓ | ✓ | ✓ | ✓ | ✓ | This work |
| SRS | Biomass composition and concentration | ✓ | ✓ | ✗ | ✓ | ✓ | ✓ | ✓ | ✓ | ✓ | 25,29 |
| SMR | Buoyant mass, volume, density | ✓ | ✓ | ✓ | ✗ | ✗ | ✓ | ✗ | ✗ | ✓ | 12–14,21 |
| Quantitative phase imaging | Refractive index (dry mass density) | ✓ | ✓ | ✓ | ✓ | ✗ | ✓ | ✓ | ✗ | ✓ | 15–17,23,24 |
| Particle tracking, FCS, FRAP | Diffusion | ✗ | ✓ | ✗ | ✓ | ✗ | ✓ | ✓ | ✓ | ✓ | 9,30 |
| Inertial picobalance | Buoyant mass | ✓ | ✗ | ✓ | ✗ | ✗ | ✗ | ✓ | ✗ | ✓ | 18 |
| Electron microscopy | Number and volume of cellular structures | ✓ | ✓ | ✗ | ✓ | ✗ | ✓ | ✓ | ✓ | ✗ | 8,9 |
| Light microscopy | Protein or lipid stain, volume of cell or nucleus | ✓ | ✓ | ✗ | ✓ | ✓ | ✓ | ✓ | ✓ | ✓ | 4,10,11,31,32 |
| Flow cytometry | Light scattering, protein stain, UV absorption | ✓ | ✗ | ✗ | ✗ | ✓ | ✓ | ✗ | ✗ | ✓ | 19,20 |

\* Absolute quantification of mass or mass density in physical units. Volume, area, or intensity in arbitrary units are considered semi-quantitative or relative measurements of biomass.

NoRI provides two important advantages over existing methods of single cell mass measurement: Protein mass, lipid mass, and water content can be measured separately, and in tissue samples with 3-dimensional resolution (Supplementary Fig. 4). The absolute calibration enables cross-comparison across contexts, and enables further specification of features based on the values without internal reference. Using NoRI, we demonstrate that protein and lipid concentration are reproducible cellular phenotypes, that they are specific to cell types, and that they change with physiological state. NoRI will enable previously inaccessible single cell mass measurements, and will allow many open biological questions to be addressed.

## Principle of Normalized Raman Imaging (NoRI)

We visualize proteins and lipids under a custom-built SRS microscope (Fig. 1a) by imaging the Raman peaks at 2935 and 2853 cm^-1^ originating from methyl groups and methylene groups^29^. For the normalization of light-scattering, SRS signals from water was measured at the 3420 cm^-1^ peak of the oxygen-hydrogen stretching modes (Fig. 1c). These three Raman bands are referred to as the CH_3_ band, the CH_2_ band and the H_2_O band in the rest of this manuscript (Fig. 1b). The different SRS intensities of proteins, lipids and water at these bands enable spectral decomposition to protein, lipid and water^33^. The mapping from the CH_2_, CH_3_ and H_2_O SRS intensity to the protein, lipid, and water concentration can be expressed with a 3×3 matrix which we refer to as the decomposition matrix. The concentration computed by conventional methods of spectral decomposition is affected by light scattering that varies within the sample and between different samples, and limits quantitative interpretation. To overcome this limitation, we took advantage of the fact that water, proteins, and lipids account for nearly 100% of the chemical components of many biological samples^34^. Hence, we scale the intensity of each component to make the sum of their volumes 1 (which is 100% v/ v) at each voxel. This results in the absolute concentration of protein, lipid and water in the unit of volume fractions. As an example, Fig. 1d shows the x-y and orthogonal cross sections of a live A7 cell’s SRS intensity images pre-normalization. Especially, the shadows in the orthogonal view of H_2_O band highlights the attenuation due to diffraction at the cell edge and intracellular lipid droplets. By contrast, NoRI normalization eliminates these variations and provides the absolute concentration of protein, lipid and water (Fig. 1e and Supplementary Fig. 18).

**Figure 1.**
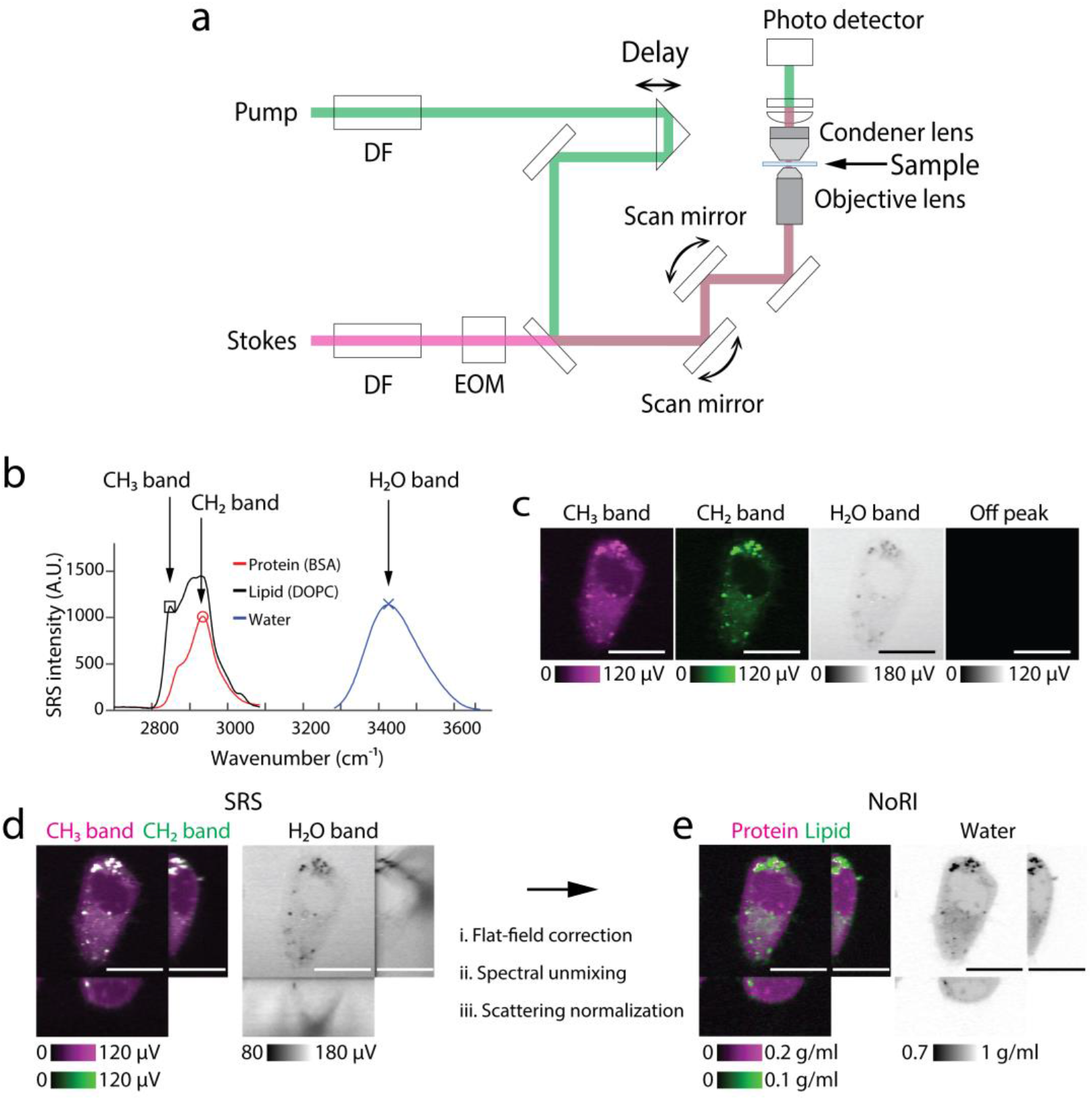
Principle of Normalized Raman Imaging (NoRI) (a) Schematic of a spectral-focusing stimulated Raman scattering microscope. Pump and Stokes femto-second pulse lasers are chirped by dense flint (DF) glass and intensity modulated by an electro-optical modulator (EOM). Motorized delay is used to fine-tune the Raman band. Transmitted pump laser intensity is detected in the trans-side. (b) Raman spectra of protein, lipid and water measured by spectral-focusing stimulated Raman scattering microscope. See Supplementary Methods for the acquisition parameters. Three Raman bands are measured in NoRI: CH_3_ (Red circle, 2935 cm^-1^), CH_2_ (Black square, 2853 cm^-1^) and H_2_O (Blue X, 3420 cm^-1^) bands. (c) Representative SRS images of a live A7 cell at the CH_3_, CH_2_, H_2_O and off-peak Raman bands. Scale bar, 20 μm. (d) Orthogonal view of the SRS images in the CH_3_, CH_2_ and H_2_O Raman bands. Scale bar, 20 μm. (e) Orthogonal view of the NoRI measurement of protein, lipid and water concentrations. Scale bar, 20 μm.

To enable this normalization, we devised a calibration and sample imaging procedure that preserves the quantitative relation between the SRS intensity and concentration (in v/v) by eliminating other sources of intensity variation. Specifically, the decomposition matrix is measured from the calibration standard samples of known protein, lipid and water concentrations using the SRS microscope (Supplementary Fig. 5). We used bovine serum albumin (BSA) as a protein standard and dioleoylphosphatidylcholine (DOPC) as a lipid standard. Since BSA has similar methyl and methylene group frequency to vertebrate proteomes (Supplementary Fig. 16), and DOPC is the most abundant lipid species in lipid membranes, they provide a practical and economical approximation of average proteome and phospholipids. To capture the intrinsic difference in Raman spectra between protein, lipid, and water samples, we measured them under the identical optical condition. The calibration images were acquired with the objective lens and the condenser lens at the same positions and the intensity variation caused by the pointing direction of the laser beams were computationally corrected from the 2D intensity profile at each tunable-laser wavelength (“flat-field correction mask”). (See Supplementary Methods for details) Once the decomposition matrix and flat-field correction mask are determined from the calibration samples, biological samples such as live or fixed cells, tissue slices and small organisms in unstained or fluorescence-labeled state can be measured. Once the raw SRS intensity images are acquired at the CH_3_, CH_2_ and H_2_O bands (Fig. 1d), the flat-field correction mask generated in the calibration step is applied to the sample images, then the decomposition matrix is multiplied to obtain the un-normalized image of protein, lipid, and water components. Then, normalization is performed at each voxel by dividing by the sum of the pre-normalization protein, lipid, and water images (Supplementary Fig. 7). The mass concentrations of proteins and lipids is estimated from the volume concentration by multiplying with the mass density of pure protein or lipids (Fig. 1e). We used the densities of BSA and DOPC (1.3643 g/ml and 1.0101 g/ml) respectively for the mass concentration conversion. We note that lipid droplets, which are composed of neutral lipids and cholesterol esters, have a shifted CH_2_ band compared to phospholipids, and, therefore, spectral decomposition based on DOPC spectrum is not ideal. On the other hand, lipid droplets occupy a distinct non-aqueous phase and the NoRI output provides an easy means to segment lipid droplets from the rest of cytoplasm by thresholding of high lipid concentration (Fig. 2b, Supplementary Fig. 10). Therefore, we distinguish lipid droplets from phospholipids by thresholding and converted lipid droplet’s volume to mass using the density of glyceryl trilinoleate (0.925 g/ml).

**Figure 2.**
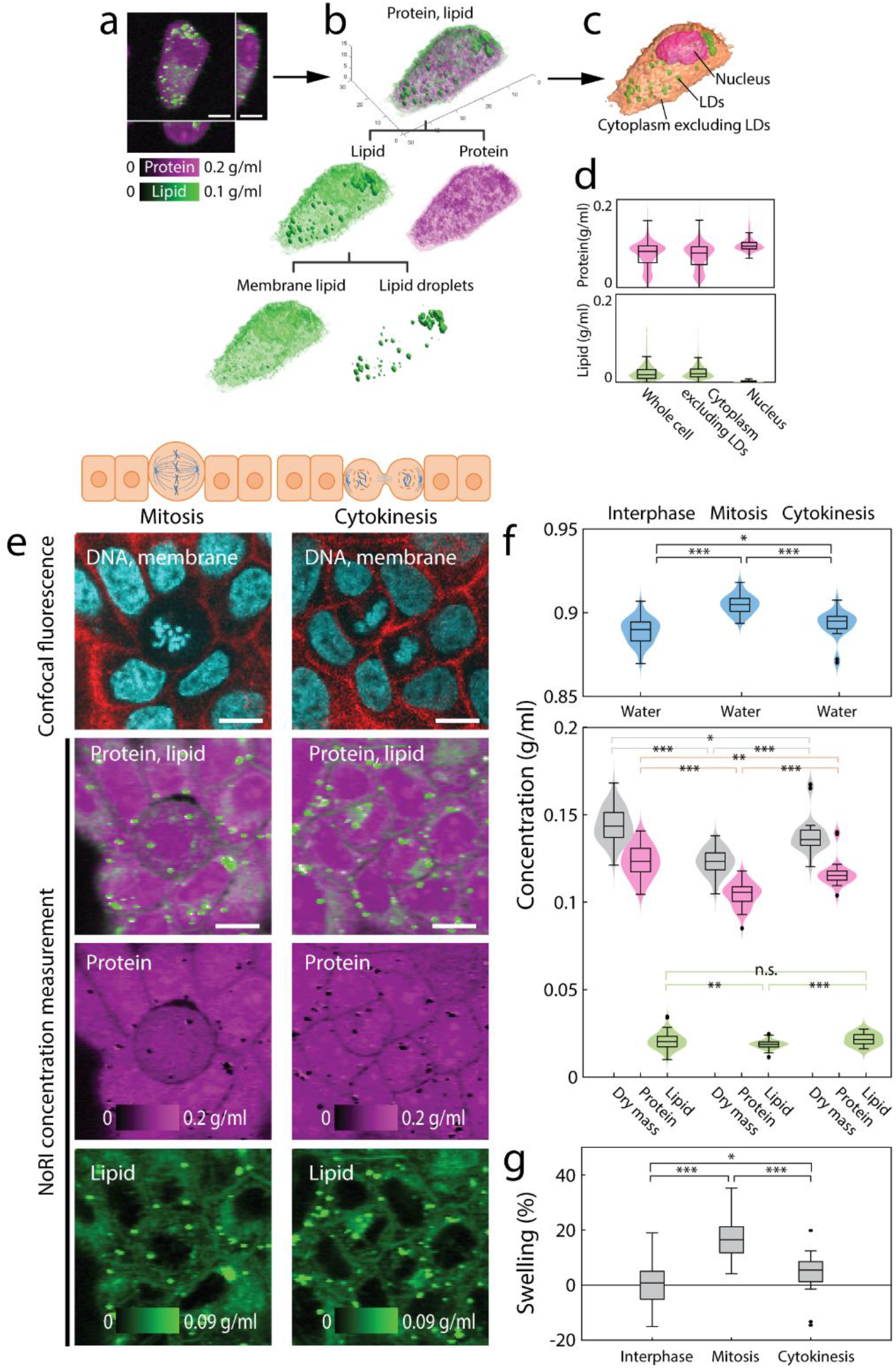
Label-free measurement of protein and lipid shows dilution of cytoplasm in mitotic cells. (a) Volumetric NoRI measurement of a live A7 cell. Scale bar, 10 μm. (b) Volumetric visualization of the protein and lipid distribution in the cell. Lipid is further classified to membrane lipids in the aqueous phase of the cytosol and lipid droplets. (c) Label-free volumetric segmentation of whole cell, nucleus and lipid droplets from NoRI data. See Supplementary Methods for the segmentation method. (d) Distribution of protein and lipid concentration within the volume of the whole cell, cytoplasm excluding lipid droplets and the nucleus in (c). (e) Representative live MDCK cells in mitosis and cytokinesis. Scale bar, 10 μm. (f) Mitotic cells have more diluted cytoplasm compared to interphase cells as revealed by lower protein and lipid concentration and higher water concentration (Water: Interphase 0.889±0.008 g/ml, mitosis 0.905±0.006 g/ml, p<0.0001. Dry mass: Interphase 0.144±0.009 g/ml, mitosis 0.123±0.007 g/ml, p<0.0001. Protein: Interphase 0.123±0.009 g/ml, mitosis 0.105±0.007 g/ml, p<0.0001. Lipid: Interphase 0.021±0.005 g/ml, mitosis 0.019±0.002 g/ml, p=0.0052). Cytoplasmic concentration recovers towards the interphase value during cytokinesis (water 0.893±0.008 g/ml, dry mass 0.138±0.010 g/ml, protein 0.117±0.008 g/ml, lipid 0.022±0.003 g/ml). Lipid droplets are excluded from concentration calculation. Number of cells in interphase, metaphase, and cytokinesis are N=81, N=57, and N=24, respectively. (g) Relative difference of cytoplasmic density is expressed in terms of the relative volume change (‘swelling’) that a cell of interphase cells’ median cytoplasmic density would undergo to manifest the observed difference in cytoplasmic water concentration. Mean and standard deviation of swelling are 0.5±6.8%, 17.0±6.8% and 4.2±7.3% for interphase, mitosis and cytokinesis respectively (p<0.001 for interphase and mitosis, p=0.020 for interphase and cytokinesis).

Next, we applied NoRI measurement to solution samples of known concentration to characterize the method’s accuracy in presence of signal attenuation of diverse origins including optical aberration from refractive index mismatching and imperfections and temporal instability of the optical system. Objective lenses are typically designed to optimally perform with an immersion media of a specific refractive index, and refractive mismatch of the sample causes optical aberration. For example, SRS intensity of 36% BSA solution sandwiched between a cover glass and a glass slide decreases with imaging depth because the BSA solution has a much higher refractive index than the intended immersion medium of the objective lens (Supplementary Fig. 8a). NoRI normalization corrects for this effect and the resulting concentration is homogenous in the entire sample volume (Supplementary Fig. 8b). Intensity variation caused by imperfections of the optical system was also removed by the normalization as demonstrated in the water component image of pure water sample (Supplementary Figs. 8c-d). Environmental instability including ambient temperature drift may cause fluctuations in SRS intensity, thereby hampering time dependent measurement. As shown in the time trace of SRS signals of a BSA solution at the CH_3_, CH_2_ and H_2_O bands, the intensity fluctuated at about 2% of the mean during 2 hours (Supplementary Figs. 8e-f). A conventional approach for normalizing would divide the CH_3_ and CH_2_ band signals with that of the H_2_O band, which indeed reduced the temporal fluctuations (Supplementary Fig. 8g). But dividing by the water signal renders the relation between intensity signal and analyte concentration nonlinear (for example, the normalization would diverge when there is little water, as in lipid droplets). NoRI normalization removes the temporal fluctuation while retaining linearity in concentration measurement (Supplementary Figs. 8h). We imaged a titration series of BSA solutions and DOPC solutions to demonstrate the linearity of NoRI concentration measurement (Supplementary Fig. 9). The NoRI measurements show excellent agreement with the actual concentration of solutions, with sensitivity of approximately 15 mg/ml as measured by the standard deviation of BSA solution sample. Since there is no existing technique to benchmark *in situ* measurement of the protein and lipid mass of a single cell, we compared the dry mass (sum of protein and lipid concentration) measurement by NoRI with refractive index tomography (Tomocube, HT-2)^35^, which is currently the only other method that can measure local mass concentration with subcellular resolution. We measured fixed HeLa cells by both techniques and found that dry mass concentration measured by NoRI was in good agreement with the dry mass concentration from refractive index tomography (Pearson correlation coefficient r=0.736 from Supplementary Fig. 12).

## Cellular protein and lipid concentration changes with cell’s physiological state

By providing protein, lipid, and water concentration measurement of a cell or a subcellular compartment, NoRI opens up new possibilities in the investigation of single cell physiology. Also, the mass of a cell or nucleus can be computed by integration of the protein and lipid concentration over the volume. Furthermore, nucleus and lipid droplets can be segmented from NoRI images without the use of staining due to their specific lipid concentration profile (Fig. 2a): The nucleus can be recognized from the absence of lipid and lipid droplets from their high lipid concentration (Fig. 2b and Supplementary Fig. 13). Here we demonstrated some potential applications.

Cells tightly maintain cytoplasmic concentration as it is directly coupled to cell volume and impacts macromolecular crowding^7,36^, structural integrity of tissue and is even reported to impact stem cell fate^37^. Though the feedback mechanism for maintaining cytoplasmic concentration against external osmotic perturbation is relatively well understood^38^, little is known about autonomous change of cytoplasmic concentration under physiological conditions^7^. Here we demonstrate that NoRI is ideally suited to probe physiologically controlled cytoplasmic concentration changes. An example of cell autonomous change of cytoplasmic concentration occurs during mitosis^39^ (Fig. 2e) concurrently with mitotic cell rounding which is important for accurate chromosome segregation^40^ and is driven by the force of osmotic pressure and actomyosin cortex contraction^41,42^. Prior reports of cytoplasmic concentration change during mitosis required two separate measurements of single cell’s total dry mass and volume and divided the mass with the volume to evaluate cytoplasmic concentration. This limited the types of sample to cell suspension or sparsely plated monolayer of cells. By contrast, NoRI enables direct measurement of cytoplasmic concentration from much more flexible sample preparations and provides additional information by distinguishing proteins and lipids. To demonstrate this, we measured the cytoplasmic protein, lipid and dry mass concentration of dividing MDCK cells in a confluent culture (Fig. 2e). The coefficient of variation of protein concentration or dry mass concentration was less than 0.07 showing that cytoplasmic concentration is tightly controlled. Also, in agreement with previous reports, the cytoplasmic dry mass concentrations decreased from interphase to mitosis and recovered after cytokinesis (Fig. 2f). The average concentration reduction during mitosis was equivalent to 17% volume swelling relative to the interphase size (Fig. 2g) and was comparable to previous reports^22,43^.

Another factor that can influence the cytoplasmic concentration is substrate stiffness. Typically, mammalian cells cultured on a stiff substrate spread to a wider area, take a flatter shape, and have higher cortical tension compared to cells cultured on a softer substrate. Even with a larger area, cells on a stiff substrate may have a lower cell volume due to the smaller height and the change in cell volume may be disproportional to protein content, leading to changes in cytoplasmic concentration^37,44^. Prior studies postulated that cells may squeeze out water upon the mechanical stimuli of the substrate through an unknown mechanism mediated by the YAP/TAZ pathway^44^. Using NoRI, we could also demonstrate similar phenomena in A7 cells cultured on polyacrylamide gels of different stiffness (Fig. 3a). A7 cells on stiffer substrate had larger area (Fig. 3b) and smaller volume (Fig. 3c), increased in water concentration (Fig. 3d) and decreased in cytoplasmic concentration of dry mass, protein, and lipid (Fig. 3e-g).

**Figure 3.**
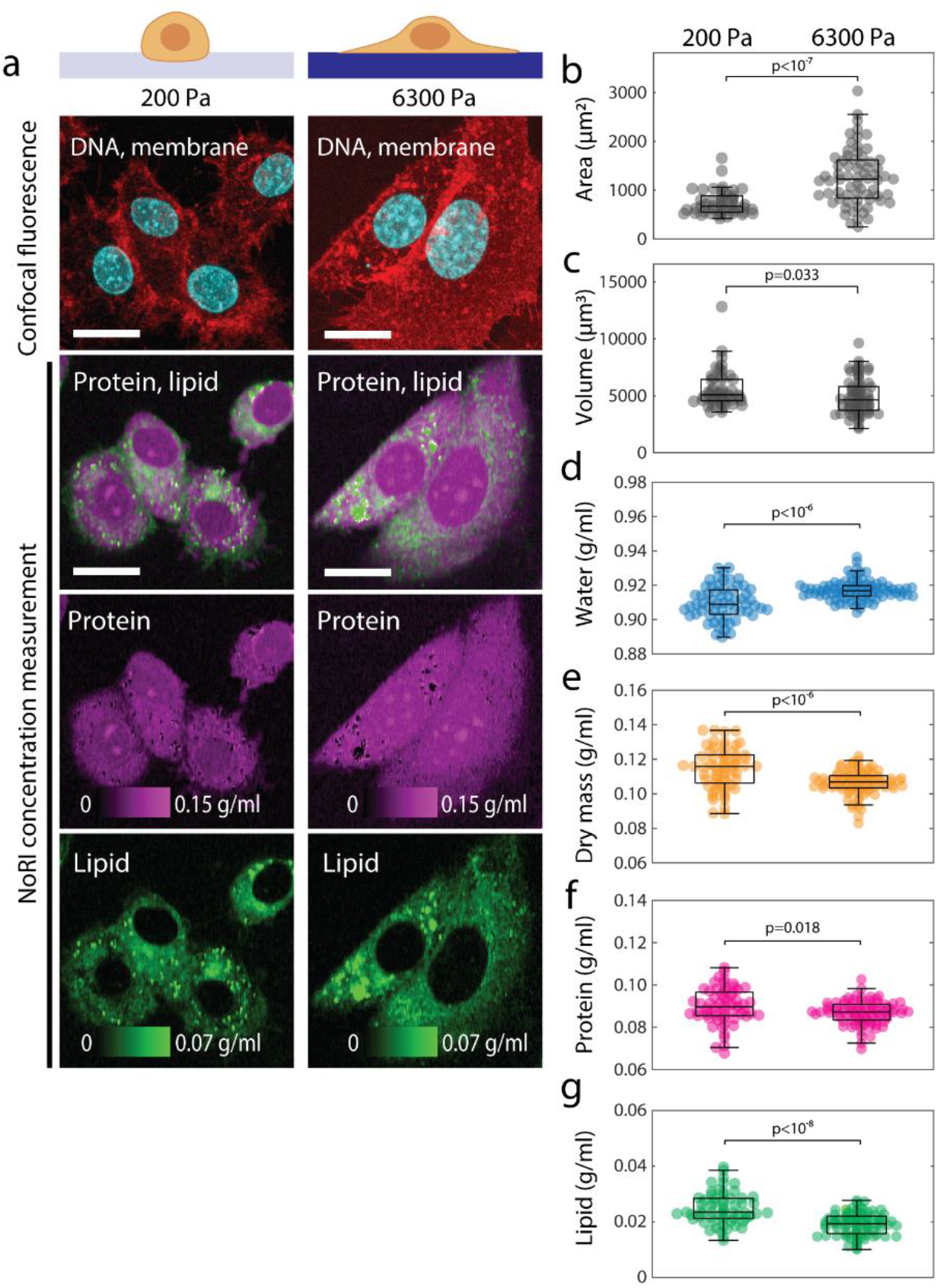
Cytoplasmic density is increased in cells cultured on a soft substrate. (a) Representative images of live A7 cells cultured for 24 hours on 200 Pa or 6300 Pa polyacrylamide gels. Scale bar, 20 μm. (b) Cell area is greater on the stiff substrate (p<10^-7^). Mean and standard deviations are 740±250 μm^2^ (N=46) and 1264±571 μm^2^ (N=67) for 200 Pa and 6300 Pa, respectively. (c) Cell volume is smaller on the stiff substrate (p=0.033). Mean and standard deviations are 5588±I672 μm^2^ (N=46) and 4921±1576 μm^2^ (N=67) for 200 Pa and 6300 Pa, respectively. (d-f) Concentration of water, dry mass (equal to the sum of protein and lipid), protein, and lipid are quantified in the single cell cytoplasm excluding lipid droplets. (d) Cytoplasmic water concentration is higher on the stiff substrate (p<10^-7^). Mean and standard deviations are 0.9098±0.0098 g/ml (N=61) and 0.9171±0.0058 g/ml (N=84) for 200 Pa and 6300 Pa, respectively. (e) Cytoplasmic dry mass concentration is lower on the stiff substrate (p<10^-6^). Mean and standard deviations are 0.1147±0.0118 g/ml (N=61) and 0.1065±0.0070 g/ml (N=84) for 200 Pa and 6300 Pa, respectively. (f) Cytoplasmic protein concentration is lower on the stiff substrate (p=0.018). Mean and standard deviations are 0.0899±0.0085 g/ml (N=61) and 0.0869±0.0055 g/ml (N=84) for 200 Pa and 6300 Pa, respectively. (g) Cytoplasmic membrane lipid concentration is lower on the stiff substrate (p<10^-8^). Mean and standard deviations are 0.0248±0.0057 g/ml (N=61) and 0.0196±0.0040 g/ml (N=84) for 200 Pa and 6300 Pa, respectively.

Next, we measured cytoplasmic concentration change during cellular senescence. Cellular senescence involves wide array of changes including hypertrophy^45^, significant cytoplasm dilution^46,47^ and accumulation of lipids^48^. Especially, dilution of cytoplasm is a poorly understood phenomena that might have a crucial role in cellular aging^46^. It was previously studied using SMR, diffusion kinetics imaging and ultracentrifugation, but these methods are either constrained to suspended cells or non-physiological preparation conditions, and lack the resolution to discern distinct processes such as lipid accumulation. NoRI provides a flexible way to observe cytoplasm dilution and lipid accumulation in senescence. To demonstrate this, we performed NoRI imaging of live MDCK cells and A7 cells induced to undergo senescence by 48 hour doxorubicin treatment (100 ng/mL) (Fig. 4, Supplementary Fig. 19). Both MDCK cells and A7 cells showed reduced cytoplasmic dry mass and protein concentrations in senescence compared to proliferating interphase cells (Fig. 4c-d, Supplementary Fig. 19c, e) in agreement with the prior report. Despite the lower dry mass concentrations, senescent cells achieve significant net hypertrophy due to cell volume increase (Fig. 4e, Supplementary Fig. 19f). We found that lipid accumulation was a significant portion of net hypertrophy in MDCK cells: The total lipid concentration (Fig. 4d), the total lipid mass (Fig. 4e), the proportion of lipid to protein (Fig. 4f), and both of the respective concentrations of lipid droplets and cytosolic membrane lipid increased (Fig. 4g) with senescence. But this disproportionate lipid accumulation was cell type dependent as senescent A7 cells showed balanced accumulation of protein and lipid (Supplementary Fig. 19f), revealing heterogeneity in metabolic reprogramming of cellular senescence.

**Figure 4.**
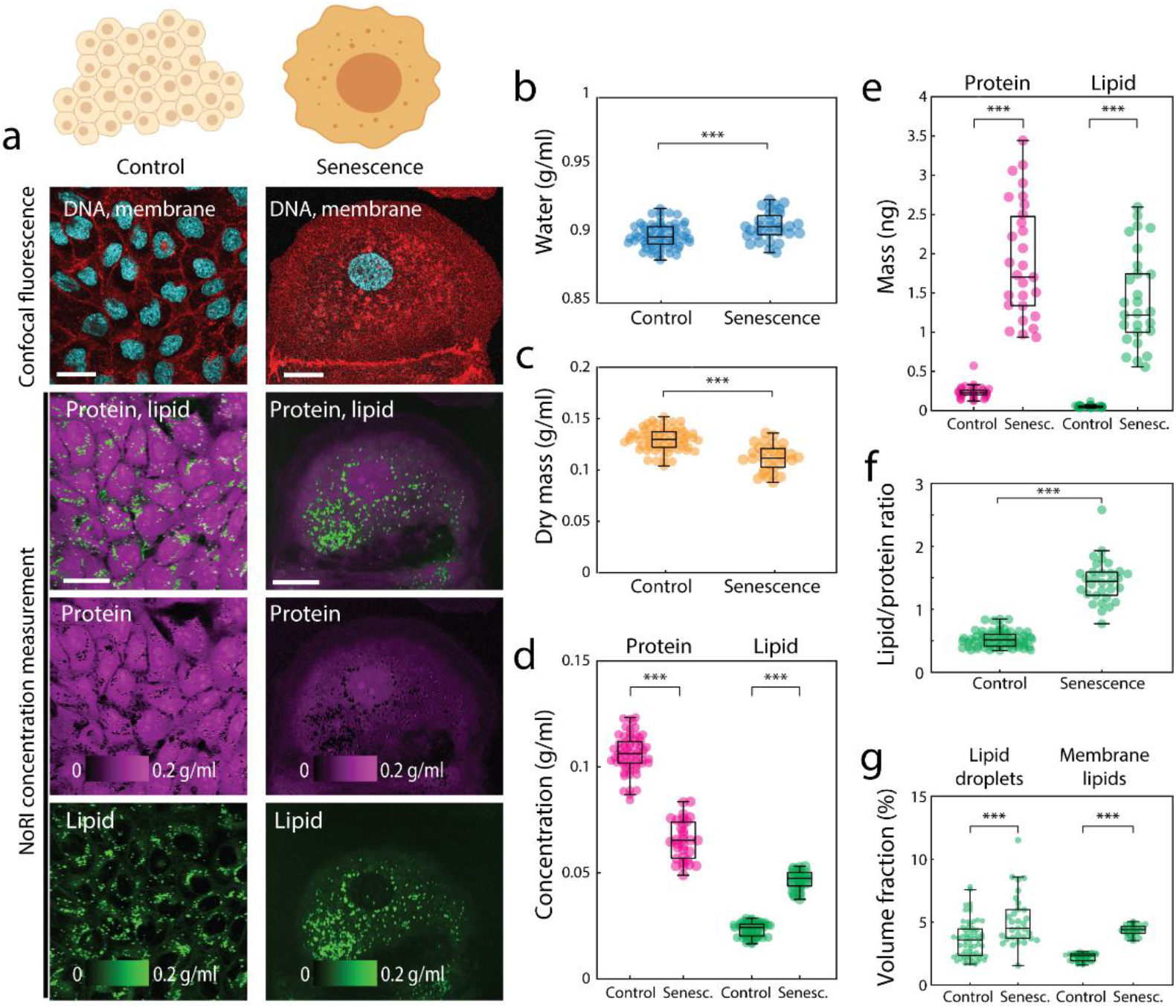
Cytoplasm dilution and lipid accumulation in senescent cells. (a) Representative images of control and senescent MDCK cells. Senescence is induced by 48 hour doxorubicin treatment. Scale bar, 30 μm. (b-d) Concentrations of water, protein, membrane lipid and dry mass of single MDCK cells. Lipid droplets are excluded from lipid and dry mass concentrations. Dry mass is calculated by adding protein mass and lipid mass. Control N=63, senescent N=31. (b) Water concentration is increased in senescent MDCK cells (p<10^-3^). Mean and standard deviation of control is 0.899±0.009 g/ml, senescent is 0.906±0.010 g/ml. (c) Dry mass concentration is decreased in senescent MDCK cells (p<10^-9^). Mean and standard deviation of control is 0.130±0.011 g/ml, senescent is 0.112±0.013 g/ml. (d) Protein concentration is decreased (p<10^-35^) and lipid concentration is increased (p<10^-47^) in senescent MDCK cells. Mean and standard deviation of control: protein 0.106±0.009 g/ml, lipid 0.023±0.003 g/ml. Senescent: protein 0.065±0.009 g/ml, lipid 0.047±0.004 g/ml. (e) Senescent cells undergo significant hypertrophy in both protein mass (p<10^-31^) and lipid mass (p<10^-31^). Single cell protein mass and lipid mass were calculated by integrating the respective concentration over cell volume. (Protein: Control 231.9±62.7 pg, senescent 1891.0±716.3 pg. Lipid: Control 50.5±14.3 pg, senescent 1377.3±588.6 pg) (f) Ratio of lipid to protein is increased in senescent MDCK cells (p<10^-32^). Total lipid is the sum of lipid droplet and membrane lipids. Lipid droplet mass is calculated from lipid droplet volume assuming density of neutral lipid 0.9 g/ml. (Control 0.53±0.13, senescent 1.44±0.34) (g) Senescent MDCK cells contain increased lipid droplets (p<10^-4^) and membrane lipids (p<10^-47^). Control: lipid droplets 3.6±1.5% (v/v), membrane lipids 2.2±0.3% (v/v), Senescent: lipid droplets 5.2±2.1%(v/v), membrane lipids 4.4±0.4%(v/v)). Lipid droplets are detected by thresholding lipid concentration >0.1 g/ml. Membrane lipid quantity is measured by integration of lipid concentration over the cell volume excluding lipid droplets.

## Cellular protein and lipid concentration are specific to cell and tissue types *in vivo*

We noted that different cell lines display different protein and lipid concentrations even in identical culture conditions suggesting cell type specificity of these properties (Fig. 5a, b). To extend this observation in animal cell types, we acquired protein and lipid concentration profiles in a diverse array of mouse tissues. Indeed, different cell types displayed distinct protein and lipid concentrations: NoRI correctly identified the differences in lipid concentrations in slow and fast skeletal muscle fibers^49^ (Fig. 5c, d). In the kidney, tubule cells showed higher lipid concentration, presumably due to densely packed mitochondria^50^, compared to glomeruli (Fig. 5e, f). Some intracellular organelles showed distinct protein and lipid concentration profile compared to the rest of the cytoplasm as in the lipid-free cell nuclei or the protein-dense zymogen granules of pancreatic acinar cells (Fig. 5g-j). The distinct protein and lipid concentration of different cell types and organelles enables interpretation of tissue microanatomy in a way comparable to the conventional hematoxylin-eosin stained histology. Furthermore, the protein and lipid concentration can provide novel information about the physiological or pathological state of the tissue. We demonstrated the potential utility of such quantitative protein and lipid concentration measurement in the following neuroscience applications. First, the quantitative difference in cytoplasmic concentration revealed hidden heterogeneity in an established cell type, the Purkinje neurons of cerebellum. These cells, identified by GAD67 expression (Fig. 6b), showed nearly 3-fold variation in the cytoplasmic concentration (Fig. 6c-d), in which lipid and protein concentrations changed proportionately (Fig. 6c). This variability was reproducible in different mouse strains (Supplementary Fig. 21b). To rule out the possibility of a fixation artefact, we performed live tissue imaging of acute brain slices and confirmed that this variability is present in live Purkinje neurons (Supplementary Fig. 21a). The difference in cytoplasmic concentration may be caused by different amount of protein synthesis or by dilution with water through an osmotic mechanism. Therefore, we sought to determine if the dense cells have more total dry mass than the light cells. For this, we acquired a volumetric NoRI image of the Purkinje layer (Fig. 6e), and traced the cell body of the Purkinje neurons (Fig. 6f). Total 12 cells were segmented from Fig. 6e and each cell’s volume, mean cytoplasmic concentration (the sum of protein and lipid concentrations) and total dry mass (the sum of protein mass and lipid mass) were calculated. We found that cytoplasmic concentration showed strong negative correlation with cell volume (Pearson’s R=-0.95), while the total dry mass shows weak correlation with cell volume (Pearson’s R=0.46). Especially, the dry mass of cells larger than 2500 fL is largely invariant with the cell volume (Pearson’s R=0.09) suggesting an osmotic mechanism of the cytoplasmic density variability.

**Figure 5.**
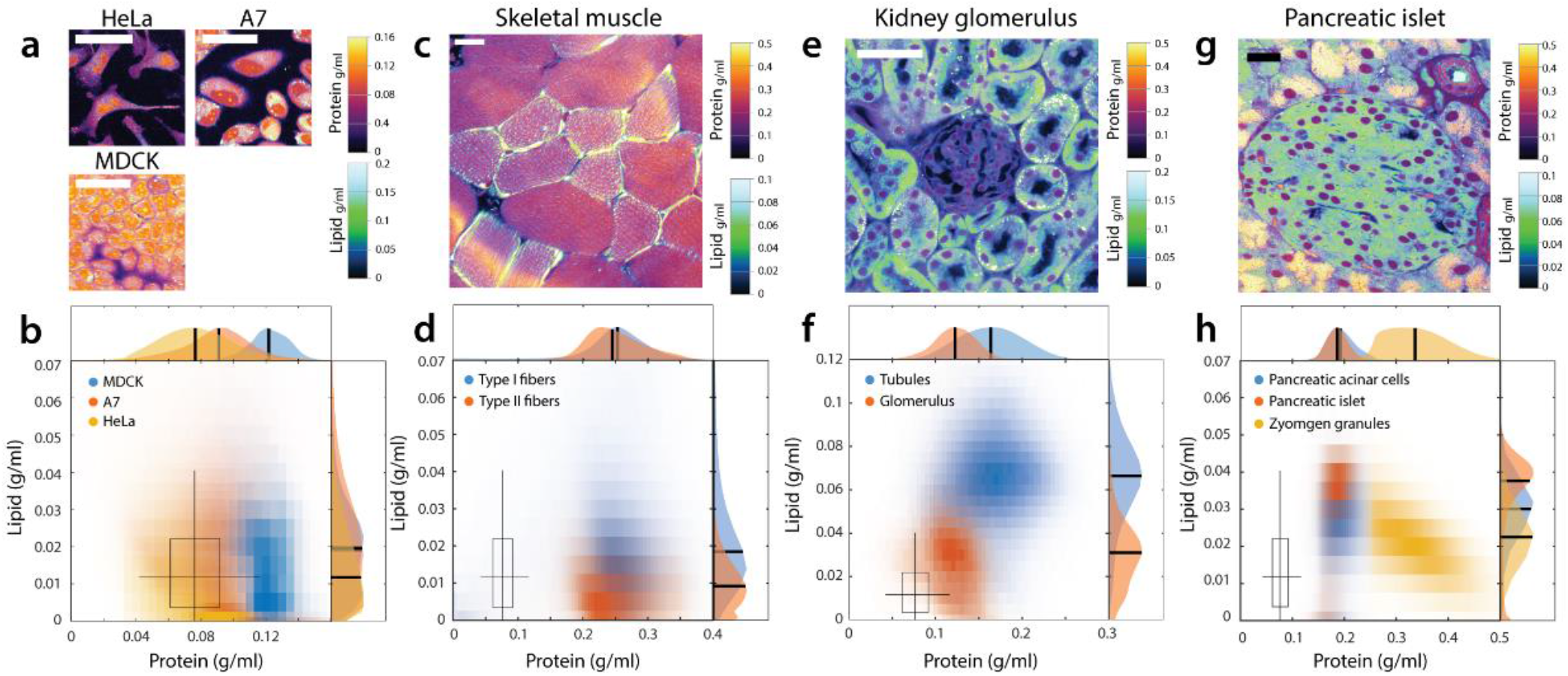
Different cell types and tissue types have distinct protein and lipid concentration. (a-b) Protein and lipid concentration distribution of fixed HeLa cells, live A7 cells and live interphase MDCK cells. Scale bar, 50μm. The boxplot marks the median, 25% and 75% quartiles of fixed HeLa cell concentrations with whiskers marking 5% and 95% range. The boxplot is reproduced in Figs. 5d, f, h, j, and n for comparison. (c-d) Protein and lipid concentration of lipid-rich Type I myofibers (N=56) and lipid-lean Type II myofibers (N=28) in traverse section of a fixed murine skeletal muscle. Scale bar, 20 μm. Type I fibers has higher lipid concentration than Type II fibers. Protein concentrations are similar between the two types. (e-f) Fixed mouse kidney tissue. Glomerulus and tubules can be distinguished by distinct protein and lipid concentration. Scale bar, 100μm. (g-h) A pancreatic islet from fixed mouse pancreas tissue. Scale bar, 100 μm. Acinar cells contained large number of protein-dense vesicles, which are likely zymogen granules for storing and secreting digestive enzymes. Cell nuclei are marked by absence of lipid. Protein dense membrane surrounds blood vessels and duct.

**Figure 6.**
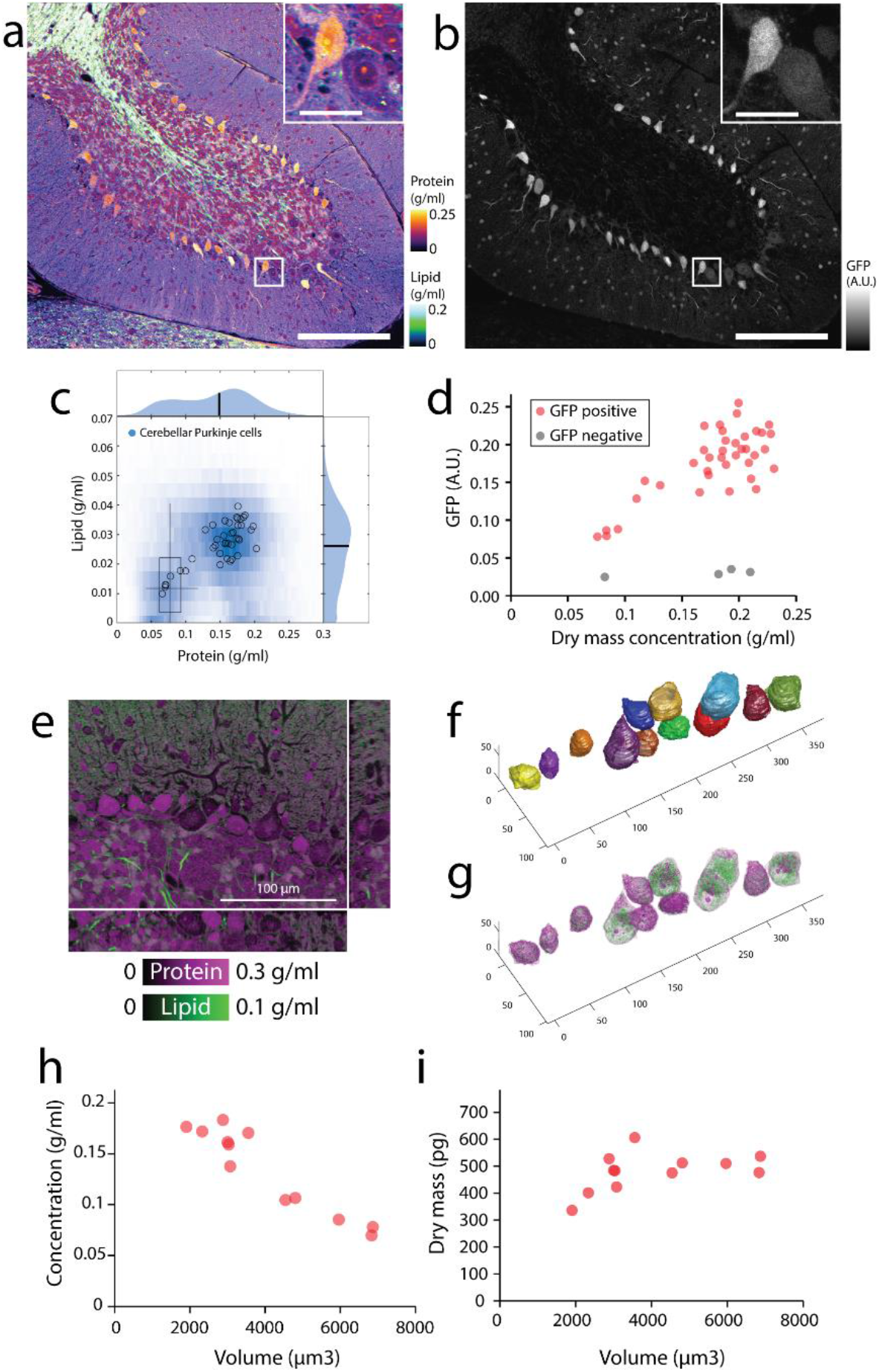
Cytoplasm density is variable in cerebellar Purkinje cells. (a) NoRI image of fixed mouse cerebellum lobe 10. Scale bar, 200μm. (Inset) Detailed view of boxed area. Inset scale bar, 30μm. (b) Confocal fluorescence of GAD67-GFP in the same area as (a). Purkinje cells are identified by expression of GAD67-GFP. (c) Protein and lipid concentration distribution of all Purkinje cells (blue density plot) and individual Purkinje cells (open circles). The boxplot displays the concentration distribution of fixed HeLa cells in Fig. 5b for comparison. (d) GAD67-GFP expression level correlates with dry mass concentration in Purkinje cells (N=38). GAD67-GFP negative cells from the Purkinje layer are also shown (N=4). (e) Volumetric NoRI measurement of fixed mouse cerebellum. Scale bar, 100 μm. (f) Cell body of Purkinje neurons are manually segmented from the label-free NoRI image in (e). Cell boundaries are determined from the distinct protein and lipid concentration of cells compared to the surrounding tissue. (g) Volumetric visualization of the protein and lipid distribution in the cells. (h) Cerebellar Purkinje cells decrease in cytoplasmic dry mass density with increasing cell body volume (N=12). (i) Dry mass of the cell body of the cells shown in (d).

## Tissue-type specific protein and lipid concentration may change with disease

Next, we explored the utility of protein and lipid measurement as a novel quantitative biomarker of disease pathology using murine models of Alzheimer’s disease^51^ (Fig. 7). SRS microscopy had been previously applied to this model and showed the capability to detect amyloid plaques by the CH_3_ Raman band^52,53^. Furthermore, it was shown that the CH_2_ Raman band can detect lipid aggregation^52,53^ in the corona of senile plaques (the area surrounding amyloid plaques^54^). In consistence with these prior reports, we observed amyloid plaques surrounded by lipid aggregates in APP-PS1 brain but not in the wild type brain (Fig. 7a, b). Since NoRI provides protein and lipid concentrations, we asked if automated pathology analysis can be performed utilizing the absolute nature of NoRI data. In this regard, we found that the protein and lipid concentrations of cell bodies, white matter, and neuropil (dense entanglement of dendrites, unmyelinated axons and synapses that surrounds cell bodies in gray matter) are similar between normal brain tissue and the normal appearing area in APP-PS1 brain tissue. The consistency of concentration profiles enabled us to perform image segmentation and detect amyloid plaques and lipid aggregates (Supplementary Methods 6.2). Briefly, cell bodies and white matter were detected by thresholding lipid images at <0.030 g/ml and at >0.147 g/ml respectively. Trainable WEKA segmentation^55^ was used to segment amyloid plaques in protein images and neuropil in lipid images. The remaining areas of intermediate lipid concentration were segmented to lipid aggregates and myelin using their morphological differences^56^. In total, we segmented the tissue into 5 classes—protein plaques, lipid aggregates, neuropil, myelin and cell body (Fig. 7c, d) utilizing their distinct concentration profiles and morphologies. The lipid aggregates were different from lipid droplets based on their much lower lipid concentration (Fig. 7e), which is consistent with the model of neurite dystrophy origin^54^. A biological replicate of APP-PS1 showed similar concentration profile in the lesions, whereas an alternative Alzheimer mouse model 5xFAD showed higher lipid concentration than those in APP-PS1 (Fig. 7f-g, Supplementary Fig. 21). Whether the quantitative difference in the concentrations signifies differences of pathophysiology in neurite dystrophy will be the subject of future work.

**Figure 7.**
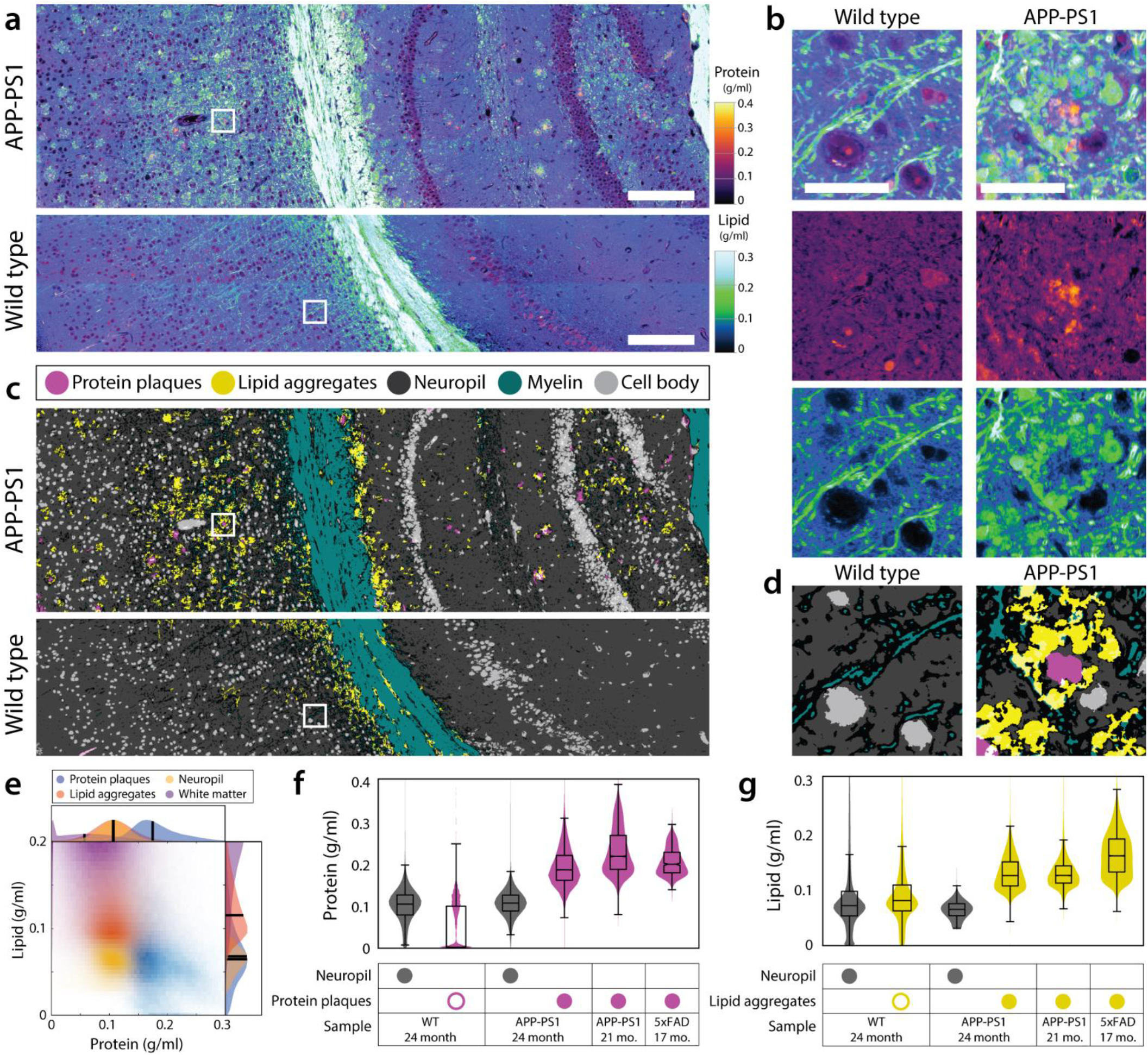
Altered protein and lipid concentration in brain tissue of Alzheimer’s disease genetic model. (a) Protein and lipid concentration images of APP-PS1 Alzheimer model mouse and wild type mouse brains. Scale bars, 100 μm. (b) Detailed view of (a). APP-PS1 brain shows senile plaques with protein-dense core and lipid-rich corona. In the wild type brain, high concentration lipid is localized to myelin fibers. Scale bar, 30 μm. (c) Tissue segmentation by using a pixel classifier based on the protein and lipid concentrations and morphological features. Both WT and APP-PS1 images are processed by the same parameters. Each pixel is classified as protein plaques (pink), lipid aggregates (yellow), neuropil (dark gray), myelin (green) or cell body (light gray). Ambiguous pixels at the border of two classes were left unclassified (black). Protein plaques and lipid aggregates are scattered throughout the gray matter of APP-PS1 mouse brain tissue. (d) Detailed view of (c). (e) Measurement of protein and lipid concentrations of protein plaques, lipid aggregates, neuropil and white matter classes in the APP-PS1 image in (a). (f) Quantitative analysis of protein concentration in WT (N=1), APP-PS1 (N=2) and 5xFAD (N=1) mice. Protein concentration is shown for neuropil and protein plaques classes detected by the pixel classifier described above. Small number of pixels in the WT were misclassified as protein plaques (open pink circle), but their protein concentration was significantly lower than that of true protein plaques (filled pink circles). The presence of protein plaques can be quantitatively characterized by the elevated protein concentration of protein plaque class. Control neuropil protein 0.099±0.042 g/ml, protein plaques 0.044±0.056 g/ml. 24 mo. APP-PS1 neuropil protein 0.106±0.031 g/ml, protein plaques 0.193±0.047 g/ml. 21 mo. APP-PS1 protein plaques 0.229±0.055 g/ml. 17 mo. 5xFAD protein plaques 0.205±0.031 g/ml. (g) Quantitative analysis of lipid concentration in the same samples as (k). Lipid concentration is shown for neuropil and lipid aggregates classes detected by the pixel classification described above. The presence of lipid aggregates can be detected by the high lipid concentration of lipid plaque class (filled yellow circle) which contrasts the low lipid concentration (open yellow circle) in the WT. Control neuropil lipid 0.089±0.071 g/ml, lipid aggregates 0.090±0.053 g/ml. 24 mo. APP-PS1 neuropil lipid 0.064±0.015 g/ml, lipid aggregates 0.129±0.030 g/ml. 21 mo. APP-PS1 lipid aggregates 0.129±0.028 g/ml. 16 mo. 5xFAD lipid aggregates.

## Discussion

Here we reported a new method, NoRI, which enables quantification of protein and lipid biomass concentrations with high spatial resolution in 3-dimensional samples. Compared to the existing methods, NoRI’s advantage is the *in situ* measurement capability and the capability to distinguish protein biomass and lipid biomass. Its working principle is based on the realization that water, protein and lipids occupy nearly 100% (v/v) of wet biological samples. Although unaccounted components such as polysaccharides limit the accuracy of NoRI measurement, the size of this error is small in typical mammalian cells where the three most abundant components (water, protein, lipid) account for ~93%(w/v) of the wet mass^34^. For example, when a sample has 75%(w/v) water, 18%(w/v) protein, and 7%(w/v) polysaccharides, the error in normalization is equal to or smaller than 7% of the calculated concentrations, i.e. <0.0525 g/ml and <0.0126 g/ml for water and proteins, respectively. Generally, water measurement permits the largest absolute error and protein and lipid measurements have small absolute error in proportion to their respective absolute concentrations. We note that the absence of water in lipid droplets does not impede the normalization even though we utilize the H_2_O band for normalization. This is because the reference is the sum of all components that is, in this case, dominated by lipid signals. Owing to its simplicity, this approach can be combined with other formats of SRS microscopy, spectral separation methods, and for the analysis of more chemical component than protein, lipids, and water. For example, the algorithm can be combined with an epi-detection SRS microscope^57^ to achieve *in vivo* measurement in live animals. Since nucleic acid displays a distinct peak in the high-wavenumber Raman band which permits spectral separation from proteins and lipids^58^, normalization method of NoRI can be used to measure the absolute concentration of nucleic acids along with protein, lipids, and water by measuring 4 Raman peaks of protein, lipids, waters and nucleic acids (Supplementary Fig. 15). To implement NoRI algorithm, quantitative reproducibility is required in the SRS instrumentation (Fig. 8). To this end we built an SRS microscope with chirped femtosecond pulse laser based on spectral-focusing principle^59^ with particular specifications to meet such reproducibility requirements (Supplementary Methods), and obtained the sensitivity of approximately 0.015 g/ml. The sensitivity can be further improved by averaging (Supplementary Fig. 14b). The spatial resolution of NoRI is determined from the resolution of the SRS microscope which is similar to that of a two-photon microscope^25^. The theoretical resolution at 770-805 nm wavelength was 0.57μm and 1.58μm in lateral and axial dimensions. Actual resolution inside tissue is usually lower due to the distortion of the laser focus by the sample, and extended in z-dimension owing to the local refractive index. The spatial resolution in tissue, as measured from the intensity profile of a small lipid droplet embedded at the 44 μm depth of a 100 μm thick tissue, was <1.18 μm and 1.90 μm in lateral and axial dimensions respectively (Supplementary Fig. 4). Pigmented samples such as red blood cells or melanocytes induce two-photon absorption (TPA) which interferes with faithful SRS intensity measurement^60^. TPA should be either measured from the off-Raman band and computationally subtracted from the Raman band signals^61^, or eliminated by bleaching the pigments^62^ (Supplementary Fig. 17). As we use light sources in 770-805 nm and 1045 nm which fall in the near-infrared window of biological tissues, dispersion is not a significant within the imaging depth (Supplementary Materials 9), yet some biological samples exhibit optical anisotropic and display different SRS intensity in a polarization sensitive way. The polarization dependence may provide additional information and is left for future works.

Both actomyosin cortex contraction and water influx are required for mitotic rounding^39,41^. Yet the mechanism of water regulation during mitosis is relatively less well understood compared to the cytoskeletal components. A few proteins including membrane proteins and ion channels had been previously identified to reduce mitotic cell swelling in an RNAi screen where cortical stiffening and volume change was measured using atomic force microscopy (AFM) and confocal microscopy^63^. Phenotypic screening using AFM is primarily aimed at mechanical properties such as stiffness and, to characterize the water flux, intracellular pressure is calculated by combining the force measurement with a biophysical model (Laplace law applied to the 3dimensional shape of the cell^39^). Cytoplasmic concentration measurement by NoRI provides a powerful alternative for probing components of intracellular water regulation. In the demonstration in this report, we observed dilution of cytoplasm in mitotic cells which was consistent with several previous reports by other groups^22,43^ (Fig. 2). However, there are also contradicting reports in which condensation of cytoplasm before or during mitosis was observed^17,64,65^. We note that the cells that swell during mitosis have interphase cytoplasm concentration between 0.095-0.190 g/ml and reduced cytoplasm concentration of 0.085-0.167 g/ml during metaphase (HT29 cells^43^, L1210 cells^22^, and MDCK cells in this study). By contrast, the cells in the latter reports, where cytoplasm condensed during mitosis, started at an extremely low interphase cytoplasm concentration of 0.025 g/ml and increased to metaphase cytoplasm concentration of 0.06 g/ml (mESC^17^). The discrepancy may be explained if the absolute cytoplasmic density rather than relative volume change is regulated during mitosis. Whether an optimal cytoplasmic density exists for mitosis is an open question which NoRI is well suited to address.

Cell shape can dramatically change depending on the stiffness of the substrate and this can be accompanied by cell volume change^44^. Cells on soft substrate usually appears more round and small while cells on stiff substrate spread to a larger area. But the spreading cells can actually have smaller volume due to reduced height^66,67^. Such cell volume changes may have functional implications such as cell fate and mediated by changes in the intracellular macromolecular crowding^37^ or membrane tension^44^. We note that A7 cells in higher stiffness gel had less dense cytoplasm as well as smaller cell volume in comparison to the cells cultured in softer gel (Fig. 3). This confirms that cytoplasm density cannot be assumed to be inversely proportional to the cell volume, which depends on the assumption of constant cellular dry mass. We believe that NoRI will be a useful tool for investigating mechanical regulation of cell volume by providing a direct measurement of the cytoplasm concentration and cell volume which is also not limited to flat substrates as existing methods^44^.

Dr. A. Amon and colleagues observed that senescent yeast cells undergo as large as 50% dilution in cytoplasm using SMR. However, in mammalian cells, the fragility of detached senescent cells prevented direct measurement using SMR and genetically encoded multimeric nanoparticles (GEMs) were used to assess the change in intracellular crowding from the diffusion rate. Several mechanisms of how cytoplasm density may impact cellular function have been proposed including dilution of unstable proteins, change of diffusion rates and reaction kinetics, and change in DNA to cytoplasm ratio^7^, but much remains unknown. By enabling direct measurement of cytoplasmic concentrations and the respective concentrations of protein and lipids, NoRI is well poised to aid further studies on the role of cytoplasm dilution in cellular senescence. Using NoRI, we confirmed that senescent mammalian cells undergo cytoplasmic dilution albeit by a smaller magnitude than yeast (Fig. 5). Along with hypertrophy, senescent cells upregulate lipid metabolism pathways and exhibit lipid droplet accumulation^45,48,68^. NoRI enables quantitative characterization of senescent cell hypertrophy and the respective contributions of protein and lipids. Using this capability, we demonstrated the heterogeneity in senescence-related lipid accumulation in different cell types. When subjected to an identical treatment and culture condition, lipid accumulation was accelerated more than protein accumulation in senescent MDCK cells (Fig. 5f), but protein and lipid accumulated proportionately in senescent A7 cells (Supplementary Fig. 19g). We believe that NoRI will be a valuable new tool for quantitative analysis of cell growth, not only in cellular senescence but in a broad range of contexts including cell cycle and development.

The difference of cytoplasmic density between cell types or cellular states has been known for a long time and was the basis for cell sorting by density gradient centrifugation^69,70^. Also, some forms of active cell volume change such as shrinking of apoptotic cells had been described^38^. However, it was only in the recent years that the development of new technologies enabled measurement of single cell cytoplasmic concentration and brought the biological regulation of cytoplasmic concentration to focus^7,36^. Changes in cytoplasmic concentration can influence protein complex formation and biochemical reaction rates via global tuning of the mass action law and diffusion rates^71^. Such global concentration tuning would have a critical effect on the activity of signaling molecules whose concentrations are close to their activation threshold^7^. Osmoregulatory components^36^, macromolecular synthesis^9^ and mechano-transduction signals^37^ are among potential factors that can modulate cytoplasmic concentrations. The functional significance of cytoplasmic concentration change and its regulatory mechanisms may be context dependent and vary with cell types: NoRI imaging of animal tissues reveals the distinct concentration profiles of different cell types (Fig. 5), heterogeneity (Fig. 6) and disease-related changes (Fig. 7). Further characterization of cytoplasmic protein and lipid concentration of various cell type in different health and disease state will be an interesting subject for future studies. Also, our observation in the Alzheimer’s disease tissue demonstrates that protein and lipid concentration can be a quantitative disease biomarker. The color intensity of histological images are usually normalized to a curve to facilitate computerbased analysis, especially to remove the color and intensity variation due to different staining protocols^72^. By contrast, absolute protein and lipid concentrations as measured by NoRI is physiologically meaningful and may provide new information. We expect NoRI to make important contributions in understanding the regulatory mechanism and functional significance of cytoplasmic concentration, as well as to open up new questions regarding the density of subcellular compartments, tissue specific problems, and questions on the coordination of lipid and protein metabolism in cell size control.

## Acknowledgements

The authors thank George Q. Daley and Ralph Weissleder for supporting the development of NoRI. S.O. and M.W.K. were supported by National Institute of General Medical Sciences (NIGMS) of the National Institute of Health (NIH) under award number R01GM026875. C.H.L. and C.J.T. were supported by NIH HD03443. W.Yang, A.L. and X.S.X. were supported by National Institute of Biomedical Imaging and Bioengineering (NIBIB) of the NIH under grant number R01EB017254. W.Yang was supported by the Office of the Provost, Faculty of Arts & Sciences, and Center for Advanced Imaging at Harvard University. The content is solely the responsibility of the authors and does not necessarily represent the official views of Harvard University. D.F. was supported by the Beckman Young Investigator Award. C.R. and W.Yin were supported by NIH R01AG055413. M.B. was supported by National Institute of General Medical Sciences (NIGMS) under award number R35GM137895. We thank Wade Regehr, Stephani Rudoph, Jui-Hsia Weng, Ahmed Rattani, Tony Tsai, Anastasia Shindyapina, José Pedro Castro, and Vadim Gladyshev for generous donation of mouse tissue samples and zebrafish embryos, Fa-ke Frank Lu, Yuanzhen Suo, Xili Liu, Doaa Megahed, Scott Gruver, Scott Rata and Victor Luria for helpful discussions, Emily Cronin-Furman, Robert Rendano and Jian Liu for technical assistance. We thank YongKeun Park and Tomocube Inc. for generously providing a demo HT-2 microscope.

## Author contributions

S.O. and C.H.L. developed the method and conducted the data acquisition and analysis. D.F. developed the initial idea of NoRI and contributed to the early experiments of the project. S.O. and A.L. constructed the SRS microscope. D.F. A.L. and W.Yang provided technical advice and manuscript revision. C.H.L. and S.O. conducted tissue imaging and single cell analysis. A.M., S.O., and C.H.L. designed and conducted *in vitro* cell culture experiments. C.R. and W.Yin provided APP-PS1 mouse model tissues and provided interpretation of the histology. C.J.T. and X.S.X. provided advice for the project and help on the manuscript. M.W.K. was involved in the conceptualization of the work and supervised the building of the microscope and the biological studies. S.O. and C.H.L and M.W.K. wrote the manuscript and all authors contributed to editing the manuscript.

## Data availability

The raw data of figures are available from the corresponding author upon request.

## Code availability

The Matlab code used in this study will be available from the corresponding author upon request.

