## Supplementary Materials for "Protein and Lipid Mass Concentration Measurement in Tissues by Stimulated Raman Scattering Microscopy"

[1]Harvard Medical School, Department of Systems Biology, 200 Longwood Avenue, Boston, MA 02115, USA
[2]Harvard Medical School, Department of Genetics, 77 Avenue Louis Pasteur, Boston, MA 02115, USA
[3]Department of Chemistry and Chemical Biology, Harvard University, Cambridge, MA 02138, USA
[4]Center for Advanced Imaging, Harvard University, Cambridge, MA 20138, USA
[5]Athinoula A. Martinos Center for Biomedical Imaging, Massachusetts General Hospital and Harvard Medical School, Boston, MA 02129, USA
[6]University of Washington, Department of Chemistry, Seattle, WA 98195 USA
[7]Beijing Advanced Innovation Center for Genomics, Peking University, Beijing, China and Biomedical Pioneering Innovation Center, Peking University, Beijing, China
[a] Co-first authors

### 1. Supplementary methods

#### 1.1. Supplementary method for the spectral-focusing stimulated Raman scattering microscope

NoRI is based on SRS microscopy of the high wavenumber Raman bands in 2853-3420 $cm^{-1}$. These Raman peaks are broad and have strong intensity and can be easily detected by many types of SRS microscope. However, not all SRS microscopes are suitable since NoRI algorithm requires that the intensity difference between different Raman bands are consistent throughout calibration and sample imaging. Use of high-stability optomechanics components, ambient temperature regulation, and encasing the free space optical components with enclosure improves repeatability. The optical system needs to be fully automated to eliminate manual optics adjustment that introduces irreproducibility between calibration and sample imaging. The most crucial element for repeatability is the light source. Tunable picosecond pulse lasers was the original choice of light source that has the advantage of straightforward Raman band selection[1]. But wavelength tuning to cover the 2853-3420 $cm^{-1}$ band required temperature change which often demanded manual adjustment of OPO cavity and Lyot filter. We found that femtosecond pulse laser (Spectra-Physics, Insight X3) has no manual adjustment and provides sufficient repeatability (Supplementary Fig. 1). It can be combined with spectral-focusing to achieve comparable spectral resolution as picosecond pulse lasers. Therefore, we built a custom SRS microscope comprised of a tunable femtosecond pulse laser and dense flint dispersion element for spectral-focusing (Fig.1a, Supplementary Fig. 2)[2].

Supplementary Fig. 2 illustrates the optical layout of the microscope. It includes a dual-output tunable femtosecond laser, dispersion elements for spectral focusing, electro-optic amplitude modulation for lock in amplification, a motorized delay for spectrum selection, a point-scanning microscope, a detector for measuring stimulated Raman loss and a lock-in amplifier. The dual-output tunable femtosecond laser (Spectra-Physics, Insight X3) provides a tunable beam (680-1040 nm) and a fixed wavelength beam (1045 nm), which are used as the pump beam and the Stokes beam, respectively. The tunable wavelength pump beam allows selection of Raman bands as depicted in Supplementary Fig. 1A. The laser power is reduced by a pair of attenuators consisting of motor controlled half wave plates and cube polarizers (Supplementary Fig. 2 A1, A2). After power attenuation, the pump beam power and the Stokes beam power at the sample were 52–88 mW and 25-39 mW, respectively. For enhanced long-term stability, beam height was reduced to 2 inches (50.8 mm) from the optical table by a periscope (not shown), and most optical components were glued to kinematic mounts affixed on 1-1.5 inch diameter optical posts made or directly mounted to the optical table. Optomechanics components made with heat-treated stainless steel with low coefficient of thermal expansion (Thorlabs, Polaris optomechanics) were used for mirror mounts, posts and post clamping arms. The pump beam and the Stokes beam are chirped by passing 4 times through high dispersion dense flint glass rods of 130 mm and 150 mm lengths, respectively (Supplementary Fig. 2 B1, B2; Casix, SF57). This chirping enhances spectral resolution and

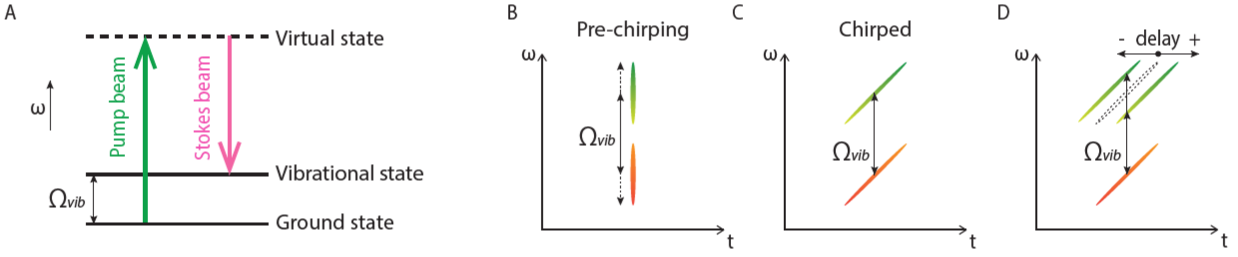


**Supplementary Figure 1. Principle of spectral focusing in stimulated Raman scattering imaging.** (A) The stimulated Raman scattering process is selective for the vibrational mode whose energy $\Omega_{vib}$ matches the energy difference between the pump beam and the Stokes beam. (B) Femtosecond pulses have a broad bandwidth, resulting in reduced resolution in vibrational spectrum. (C) Chirped pulses have narrow instantaneous bandwidth and enhances the spectral resolution. (D) The energy difference between the pump beam and the Stokes beam can be adjusted by changing the temporal overlap of the two chirped pulses. This is adjusted by a controllable delay in the optical path of the pump beam.

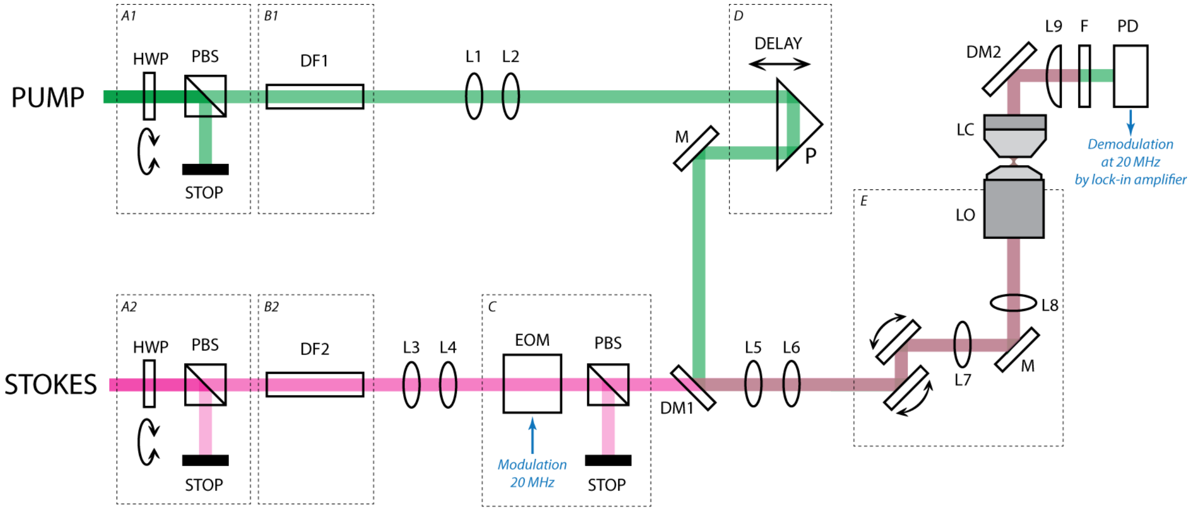


**Supplementary Figure 2. Schematics of spectral focusing SRS microscope optics.** PUMP, tunable output of the femtosecond laser; STOKES, 1045 nm output of the femtosecond laser; (A1, A2) Power attenuation modules. HWP, half waveplate; PBS, polarizing beam splitter; STOP, beam stop; (B1, B2) Pulse chirping for spectral focusing. DF1-2, SF57 glass rods; (C) Power modulation module for lock-in amplification. EOM, electro-optic modulator; (D) Motorized delay for spectrum selection and scanning. P, retroreflector prism; (E) A confocal laser scanning microscope; PD, photo diode; DM1-2, dichroic mirrors; M, mirror; L1-6, relay lens pairs for adjusting the beam diameter; L7, scan lens; L8, tube lens; LO, objective lens, LC, condenser lens.

is referred to as spectral focusing (Supplementary Fig. 1B and 1C). The amplitude of the Stokes beam is modulated by a resonant electro-optic modulator (Supplementary Fig. 2 EOM; Thorlabs EO-AM-R-20-C2) at 20 MHz (Supplementary Fig. 2 C). To drive the EOM, the sinusoidal output from a function generator internal to the lock-in amplifier (Zurich Instruments, UHFLI) is amplified by using a RF power amplifier (Mini-Circuits, ZHL-1-2W). The modulation depth is maximized by adjusting the amplitude of the driving voltage. A motor-controlled retroreflector prism controls the optical path length of the pump beam (Supplementary Fig. 2 D). This path length adjustment provides a fine control of spectrum selection (Supplementary Fig. 1D)[2]. The Stokes beam is combined with the pump beam using a dichroic mirror (Supplementary Fig. 2 DM1; Semrock, LP02-980RE-25). Path length of the pump beam path prior to the dichroic mirror is appropriately adjusted to ensure temporal overlap of the pump pulse and the Stokes pulse after the dichroic mirror. The pulse overlap can be determined at the resolution of nanoseconds (equivalent to millimeter distance of path length) using a pair of high speed photo detectors (Thorlabs, DET10A) and a high bandwidth oscilloscope (≥500 MHz). Finer adjustment of the pulse overlap is made by the motorized delay using the SRS signal from a sample. The combined beams enter a laser scanning microscope (Supplementary Fig. 2 E; Olympus, FV3000) to image the sample by point scanning. Sample is placed between a water immersion 60x 1.2NA objective lens (Supplementary Fig. 2 LO, Olympus, UPLSAPO60XWIR) and oil immersion 1.4 NA condenser (Supplementary Fig. 2 LC, Nikon, D-CUO Achr-Apl NA=1.40). The sample space is 1.83 mm when the condenser is directly immersed in the culture dish (without immersion oil) and 0.63 mm when imaging through 1.2 mm thick slide glass (with immersion oil). The transmitted light is collected by the condenser lens and focused on a high speed photodiode (Supplementary Fig. 2 PD; Thorlabs, FDS1010) located on the trans-side of the sample using a focusing lens (Supplementary Fig. 2 L9). Light path is folded by 90 degrees using a dichroic mirror (Supplementary Fig. 2 DM2; Edmund Optics, 69-218) to allow transillumination for bright field viewing of the sample through microscope eyepiece. The Stokes beam is rejected before the detector with an IR filter (Supplementary Fig. 2 F; Edmund Optics, 54-517) and stimulated Raman loss of the pump beam is measured. Bias voltage of 70 V was applied to the photodiode. The photo current is converted to voltage with a 50 Ω resister, filtered by a 3–30 MHz bandpass filter (Mini-Circuits, ZABP-16+), and demodulated by a lock-in amplifier (Zurich Instruments, UHFLI) at the EOM modulation frequency. The demodulation amplitude from the lock-in amplifier is registered to images in real time by an analog-to-digital data acquisition board (Olympus, Analog Box) synchronized to the scanning of the microscope.

The tunable laser wavelength and delay position parameters for the $CH_3$, $CH_2$, $H_2O$ and other desired Raman bands are determined as follows. The pump beam wavelength for selecting a Raman band of vibrational energy $\Omega_{vib}$ is determined by $\Omega_{vib} = 1/\lambda_p - 1/\lambda_S$, where $\lambda_p$ and $\lambda_S$ are the wavelengths of pump beam and Stokes beam, respectively (Supplementary Fig. 1A). Supplementary Table 1 shows the pump beam wavelengths for the $CH_3$, $CH_2$, $H_2O$ and nucleic acids Raman bands. Once the pump beam wavelength is set, as described above, we determine the delay position for the $CH_3$, $CH_2$, and $H_2O$ Raman peaks. For this, we acquire spectral scans of pure samples (BSA, DOPC, water, or DNA) by scanning the delay motor at the respective pump beam wavelength shown in Supplementary Table 1. Measured SRS spectrum is the convolution of the Raman spectrum and the overlap of chirped pump and Stokes pulses (Fig. 1b)[2]. The pulse overlap is a function of the delay position can be measured from two-photon absorption (TPA) of Rhodamin 6G solution (Supplementary Fig. 3a). Raw SRS intensity is divided by Rhodamin 6G TPA to obtain Raman spectrum (Supplementary Fig. 3c).

For routine imaging, the delay position at the peak of $CH_3$, $CH_2$, and $H_2O$ peaks were saved in preset files. Since the linear delay motor position and Raman spectrum wavenumber are linearly related[2], the delay position can be mapped to wavenumbers by linear fitting using the $CH_2$ and $CH_3$ Raman peaks. We wrote Matlab scripts to automate SRS image acquisition at the Raman bands. Matlab preset files stored the parameters of wavelength, delay position, and power attenuation for respective Raman bands. The setting of the scanning microscope such as the scan positions, z-scan range, and scan resolution were controlled from Olympus Fluoview, which in turn was controlled by Matlab via Remote Development Kit (Olympus, RDK) library.

**Supplementary Table 1.** Pump beam wavelengths for selected Raman bands

| Raman band | $\Omega_{vib}$ ($cm^{-1}$) | Stokes beam wavelength (nm) | Pump beam wavelength (nm) |
|---|---|---|---|
| $CH_3$ | 2935 | 1045 | 800 |
| $CH_2$ | 2853 | 1045 | 805 |
| $H_2O$ | 3420 | 1045 | 770 |
| Nucleic acids | 2967 | 1045 | 796 |

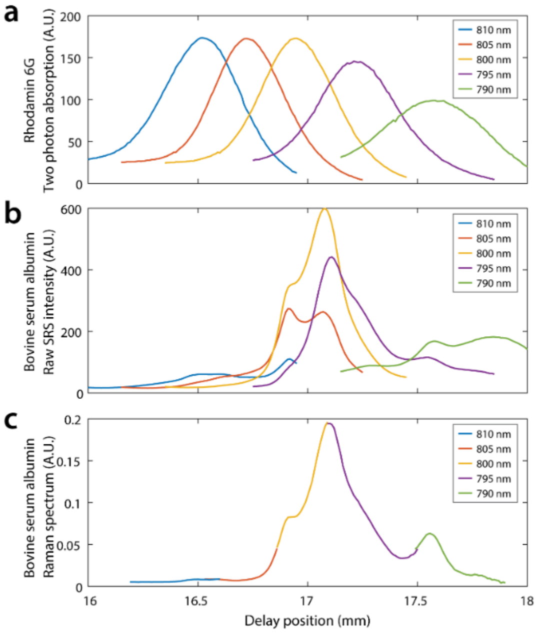


**Supplementary Figure 3. Raman spectrum measurement using spectral-focusing SRS microscope.** Spectrum is acquired by scanning the delay length of the pump beam. (a) Pulse overlap of pump beam and Stokes beam is measured from two photon absorption of Rhodamin 6G dissolved in deuterated water. Colors indicate different pump beam wavelengths as per legend. Stokes beam wavelength is 1045nm. (b) Raw SRS intensity of bovine serum albumin as a function of pump beam delay at respective pump beam wavelengths. (c) By dividing (b) with (a) after dark signal subtraction, Raman spectrum of bovine serum albumin is obtained. Data is truncated at the edges of each scan where noise is large due to low two photo absorption signal.

#### 1.2. Image acquisition and resolution

All SRS images are acquired with 2 microsecond per pixel dwell time. As such, the acquisition time for a single 512x512 resolution SRS image is about 1 second. In a typical NoRI measurement, three Raman bands are acquired at 2853, 2935, and 3250 $cm^{-1}$. A Raman band change requires a wavelength change of the tunable laser, which takes 7–11 seconds to finish and limits the rate of time series data acquisition. When scanning a large sample area of fixed sample, we imaged the entire sample area at each Raman band to minimize the time switching pump beam wavelength. In these cases, we minimized image misalignment between different Raman bands by clamping the sample to the stage and equilibrating it with the environmental temperature before scanning. At the 60x magnification, the field of view is approximately 212 μm x 212 μm. Therefore, the time to image a 1 cm x 1 cm area at a single z plane is about 2 hours. Frame averaging of 4 to 16 frames were used in 4-component NoRI imaging for nucleic acid concentration measurement (Supplementary Fig. 15). No frame averaging was used in other cases unless noted. Theoretical resolution at 60x magnification was 0.34 μm laterally and 0.85 μm axially[3] (0.57 μm and 1.58 μm when converted to full-width-half-maximum[4]). To demonstrate the practical resolution in tissue imaging, we measured a lipid droplet located at 44 μm depth inside a 100 μm thick section of cartilage tissue without frame averaging (Supplementary Fig. 4). Full-width-half-maximum of the lipid droplet was 1.18 μm and 0.714 μm in x and y and 1.90 μm in z, which is slightly larger than the theoretical resolution.

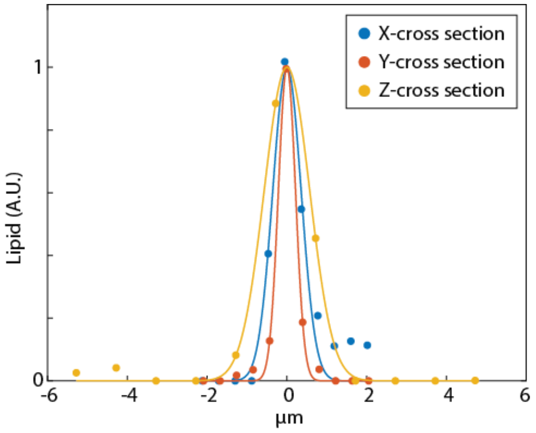


**Supplementary Figure 4. Cross section of a small lipid droplet.**

#### 1.3. Pre-processing of SRS images

To ensure the linearity of the SRS intensity in the entire signal range including small signals near zero, in-phase amplitude (X) is chosen for lock-in amplifier output and a small constant offset is added to the output. The offset is subtracted after analog to digital conversion by subtracting the background level measured when the light is blocked. After the background subtraction (which is same for all Raman bands), SRS images were divided by the flat field correction mask of the matching Raman band. This is necessary because the SRS intensity is stronger at the center of the field of view than in the periphery due to the aberration of the optical system. This spatial variation of intensity does not change with time or repeated changing of the pump beam wavelength, but are different for different pump wavelengths. Flat-field correction masks were created by measuring the intensity profile of BSA solution at 770 nm, 796 nm, and 800 nm pump wavelengths (for the $H_2O$, nucleic acids, and $CH_3$ Raman bands respectively) and from DOPC solution at 805 nm pump wavelength (for the $CH_2$ Raman band). SRS intensity image at the z plane of the highest intensity is smoothed by 2D polynomial fitting[5] and normalized by dividing with the maximum intensity. The mask is generated from a single z plane of the calibration samples but applied to all z planes in the sample data.

#### 1.4. Protocol for making calibration standards

Calibration standards are bovine serum albumin (BSA) solution in water, dioleoyl-phosphocholine (DOPC) solution in 4-deuterated methanol, pure water and pure 4-deuterated methanol. All solution samples were mixed by measuring the weights of the components. BSA powder (Sigma Aldrich, A7638) was weighed in a conical tube and 150 mM pH7.6 phosphate buffer was added to make the concentration of 30-33% (w/w). The tube is tightly closed and placed in a shaker at 40 °C until BSA is fully dissolved, while occasionally spinning down undissolved chunks stuck on the tube wall. DOPC powder (Avanti Polar Lipids, 850375) was weighed in a conical tube, to which 4-deuterated methanol (Sigma-Aldrich, 151947) was added to the final concentration of 15-39%. Double strand DNA solution was used in the 4-component NoRI (Supplementary Fig. 15). Double stranded DNA powder from salmon testes (Calbiochem, 262012) was weighted in a conical tube and water was added to make the final concentration 5% (w/v). The tube was tightly closed and placed in a shaker at 37 °C overnight until DNA was dissolved. The final concentration in volume fraction was calculated from the mass fractions and the density of the pure components, which is 1.364 g/ml for BSA[6], 1.01 g/ml for DOPC[7], and 1.7 g/ml for DNA[8].

#### 1.5. Protocol for measuring SRS intensity of the calibration standards

To compute decomposition matrix, calibration standard samples need to be imaged under the same imaging condition. This can be achieved by flowing different sample solutions to a single same imaging chamber, but this method requires that the fluidics system is flushed between samples and large quantity of standard solution is consumed in the process (Supplementary Fig. 5a). We find it convenient to assemble all calibration standard samples on a glass slide by creating multiple sample holding slots using adhesive sheets (Grace Biolabs, SA-S-1L)(Supplementary Fig. 5b). The slots were created with regular spacing using a laser cutter to automate the image acquisition. The size of the sample holding slot was 2 mm width, 24 mm length, and 120 μm thickness and held about 6 μl of solution. To prevent mixing of the adjacent samples, we added empty slots around the DOPC sample and the d-methanol sample. The samples were injected using a micropipette and sealed with vacuum grease. Supplementary Fig. 5c shows SRS intensity z-profiles of each sample measured at the $CH_3$, $CH_2$ and $H_2O$ Raman bands. The SRS intensity of the water sample is independent of the z position, as the objective lens is corrected for water. But the intensity of BSA and DOPC samples is strongest at the surface immediately past the cover glass and decreases with increasing z-depth as the focus degrades as light pass through the index-mismatched solutions. For the computation of the spectral decomposition matrix, the average SRS intensity values are measured at the z plane of highest intensity for protein and lipid, and at the mid-z-plane for water and 4-deutrated methanol. This measurement was performed after background-subtraction and flat-field-correction of the raw SRS intensity images.

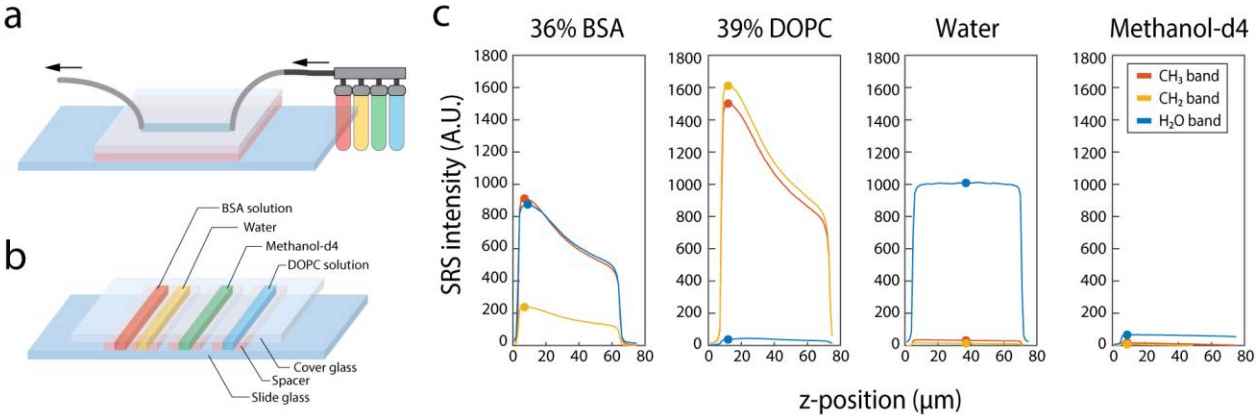


**Supplementary Figure 5.** (a) Flow channel for measuring calibration samples under an identical imaging condition. (b) Multi-channel slide assembly for calibration samples. (c) Z-profiles of SRS intensity of calibration standard samples. Filled circles mark where the SRS signal intensity is taken for the calculation of the decomposition matrix.

##### 1.6. Computation of the decomposition matrix

We obtained the spectral decomposition matrix that maps $\mathsf{S}$, the SRS intensities of the calibration samples at the $CH_3$, $CH_2$, and $H_2O$ Raman bands, to $\mathsf{C}$, the protein, lipid, and water concentration of the calibration samples as the following. The matrix $\mathsf{S}$ is defined as a 3x4 matrix where $\mathsf{S}_{ik}$ denotes SRS intensities of the BSA standard solution, water, the DOPC standard solution, and 4-deuterated methanol ($k$=1–4) at the $CH_3$, $CH_2$, and $H_2O$ Raman bands ($i$=1–3). The intensities are calculated after the background subtraction and flat field correction from the average of the whole image at the single z plane as described above (Supplementary Fig.5c). The matrix $\mathsf{C}$ is defined as a 4x4 matrix where $\mathsf{C}_{jk}$ denotes the concentration of protein, lipid, water and deuterated methanol fractions ($j$=1–4) in the standard samples ($k$=1–4). The concentration is expressed in the unit of volume fractions between 0 and 1. Decomposition matrix is computed by solving for $\mathsf{M}$ that satisfies $\mathsf{C} = \mathsf{MS}$. This can be computed in a single line using right matrix division in Matlab $\mathsf{M} = \mathsf{C}/\mathsf{S}$, which gives a 4x3 matrix that maps three Raman bands to protein, lipid, water and deuterated methanol fractions. The last row that maps to deuterated methanol is not necessary and dropped out (they are also inaccurate as the SRS signal is practically zero in all three Raman bands), resulting in a final 3x3 matrix form of $\mathsf{M}$.

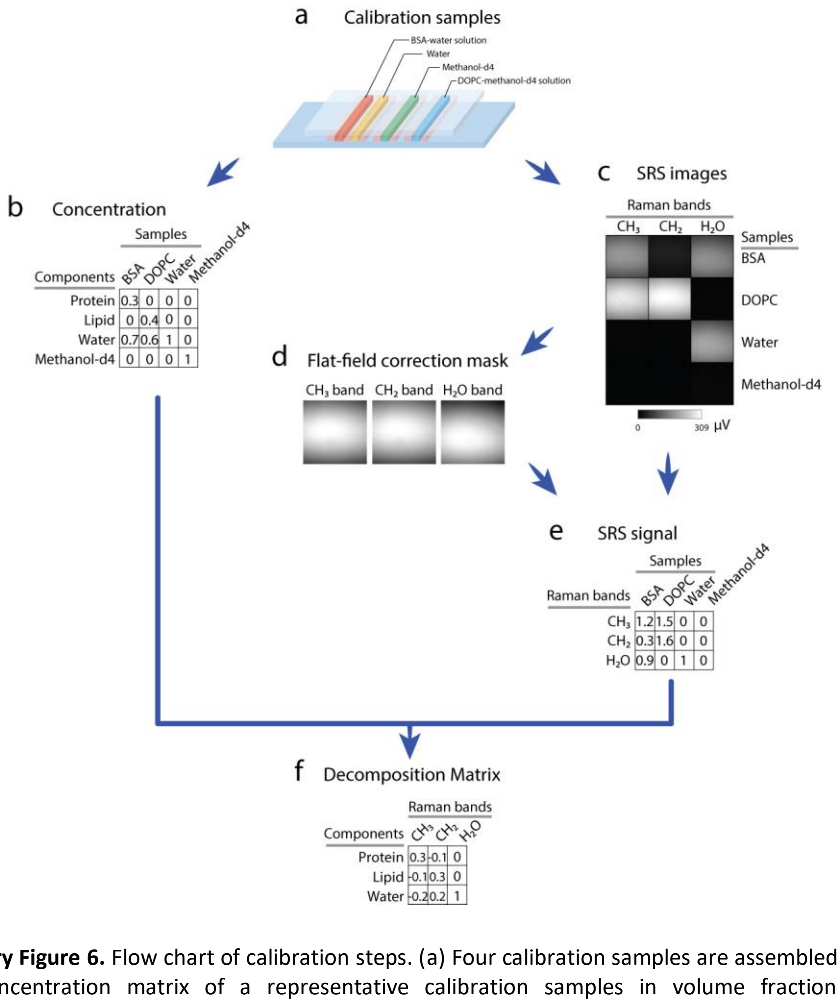


**Supplementary Figure 6.** Flow chart of calibration steps. (a) Four calibration samples are assembled on a slide glass. (b) Concentration matrix of a representative calibration samples in volume fraction unit. (c) Representative raw SRS intensity images of 4 calibration standard samples at the $CH_3$, $CH_2$, and $H_2O$ Raman bands. (d) Flat field correction masks are computed for each Raman band. (e) SRS signal matrix is calculated by taking the average of the whole image after background subtraction and flat-field correction. (f) Decomposition matrix is computed by solving for M in C=M*S where S is the SRS signal matrix and C is the concentration matrix.

##### 1.7. Applying the decomposition matrix to sample data

Decomposition matrix is multiplied to sample images after background subtraction and flat-field correction. Spectral decomposition is applied to individual pixels independently by multiplying the 3x3 decomposition matrix to the 3-component vector of the SRS intensities at the $CH_3$, $CH_2$, and $H_2O$ Raman bands of each pixel. This matrix multiplication can be simplified by linearizing the image data as a 1xN vector. The linearized images of three Raman band can be stacked as a single 3xN matrix, S. The spectral decomposition becomes matrix multiplication of a 3x3 matrix M with a 3xN matrix S. Then result is 3xN matrix where each row represents the pre-normalization concentration of respective chemical species. Each row can be reshaped to the original image size to obtain the decomposition image.

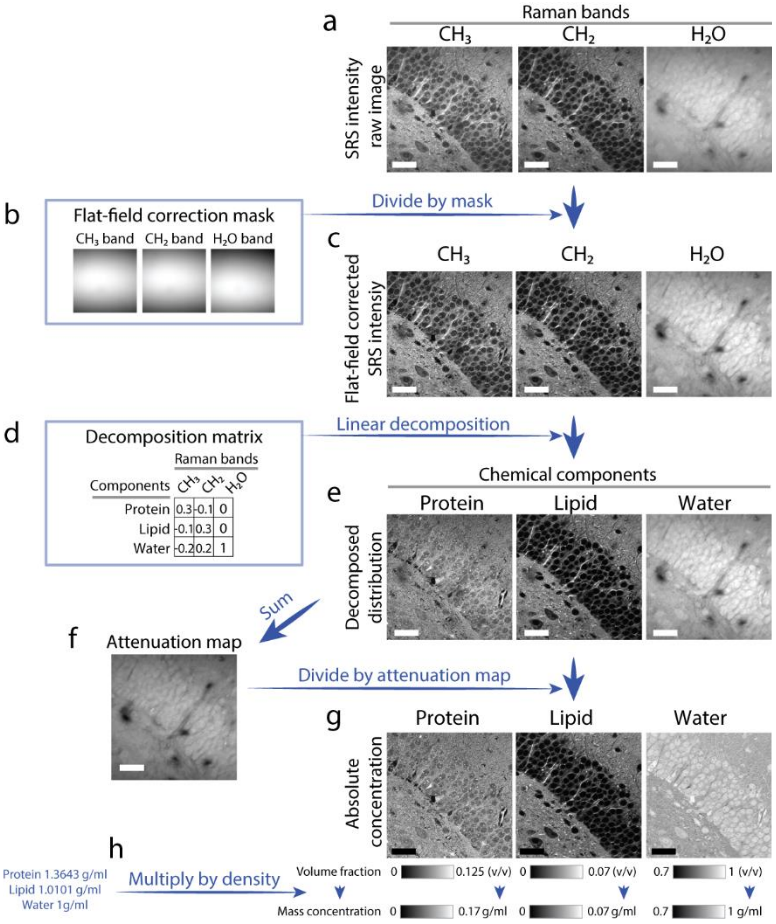


**Supplementary Figure 7. Flow chart of sample data processing steps.** (a) Sample is imaged at the $CH_3$, $CH_2$ and $H_2O$ Raman bands and the background level is subtracted. (b) Flat field masks are generated from the calibration data. (c) Sample images are divided by the flat field correction mask of the matching Raman band. (d) Decomposition matrix is acquired in the calibration step. (e) Decomposition matrix is multiplied to each pixel, generating pre-normalization concentration images. (f) The pre-normalization protein, lipid, and water images are summed at each pixel to compute the attenuation map. (g) Each pre-normalization image is divided by the sum image (attenuation map) to compute the absolute concentration image in the unit of volume fractions. (h) Volume fraction is multiplied by the density of pure BSA, DOPC, and water to convert to mass concentration.

#### 1.8. Demonstration of NoRI measurements

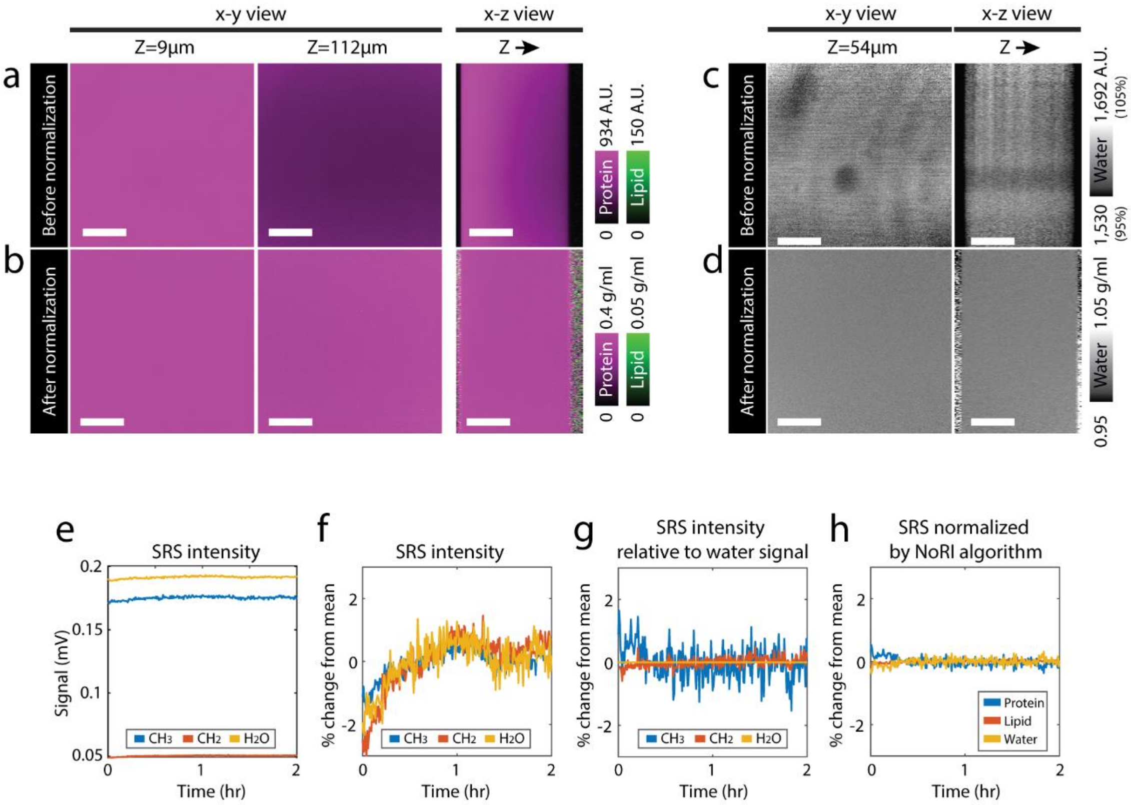


**Supplementary Figure 8.** NoRI eliminates spatial and temporal variability in measurements. (a) Pre-normalization image of protein and lipid concentration of 33%(w/w) BSA solution. The solution is sandwiched between a coverslip and a slide glass. Signal intensity diminishes with z depth due to optical aberration caused by the high refractive index of the solution. Scale bar, 50 μm. (b) NoRI normalization of (a) removes the effect of optical aberration and enables accurate measurement of concentration throughout sample volume. Scale bar, 50 μm. (c) Pre-normalization water concentration image of water sample shows stationary spatial variation caused by imperfections of the optical system and detector. Scale bar, 50 μm. (d) NoRI normalization is applied to (c) and removes the stationary spatial variation. Scale bar, 50 μm. (e) SRS intensity from a BSA solution is acquired at the CH3, CH2 and H2O bands every 30 seconds for 2 hours. (f) Data in (e) is plotted as percentage changes relative to the mean. Temporal fluctuation due to environmental and instrumental instability is about 2% and shows a similar trend in all 3 Raman bands. (g) SRS signal fluctuation relative to the $H_2O$ band signal. The $CH_3$ band and $CH_2$ bands are divided by $H_2O$ band signal at each time point. (h) NoRI normalization eliminates temporal fluctuations better than relative measurement to the $H_2O$ band signal.

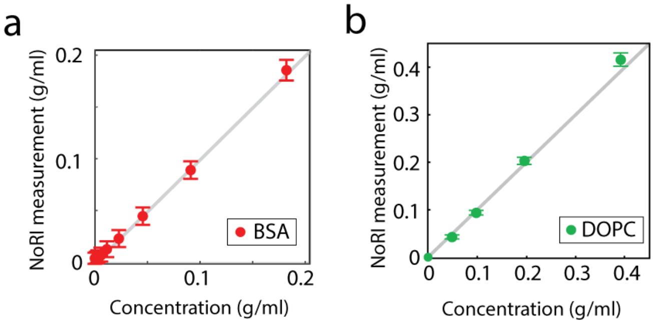


**Supplementary Figure 9.** NoRI measurement of (a) protein concentration of BSA solutions and (b) lipid concentration of DOPC solutions show good agreement with the actual concentration. In DOPC titration, organic solvent (deuterated methanol) was used in place of water for normalization. Error bars indicate standard deviations.

#### 1.9. Simultaneous fluorescence imaging with NoRI

NoRI is compatible with fluorescence imaging and simultaneous or tandem acquisition of confocal and two photon fluorescence is possible using a custom 5-band filter (Supplementary Fig. 10A-B DM2; Chroma, zt405/488/561/640/NIR-xr-rpc-UF2). An example of tandem fluorescence and NoRI imaging is shown in Supplementary Fig. 11.

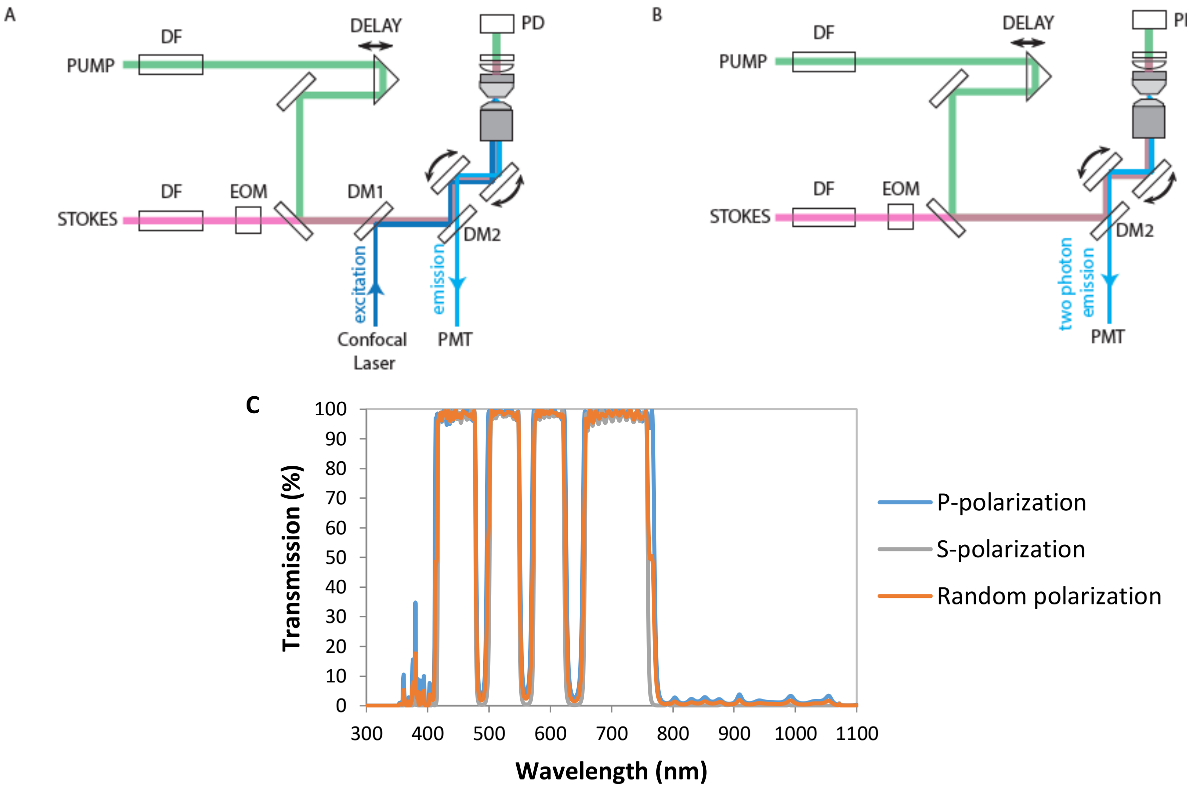


**Supplementary Figure 10. Schematics of simultaneous or tandem acquisition of fluorescence with NoRI.** (A) Light path for simultaneous confocal fluorescence and SRS imaging. DM1 is a longpass filter for combining combine confocal excitation lasers (405, 488, 561, 640 nm) with near-infrared lasers for NoRI (770 – 1045 nm). DM2 is a customized 5-band pass filter. (B) Light path for simultaneous two photon fluorescence and SRS imaging. Confocal pinhole is fully opened. (C) Transmission spectra of the custom 5-band filter.

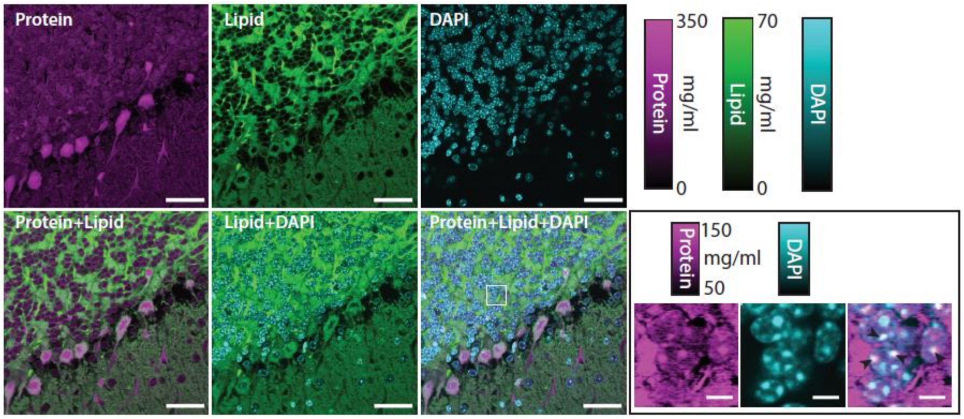


**Supplementary Figure 11. Tandem acquisition of DAPI fluorescence and NoRI.** DAPI fluorescence by confocal fluorescence and protein and lipid NoRI images of fixed mouse cerebellum. Scale bar, 40 μm. (Inset) Magnified view of DAPI and protein puncta. Overlapping DAPI and protein puncta are marked with arrowheads. Inset scale bar, 5 μm.

#### 2. Cell culture sample preparation

A7 human melanoma cell line was purchased from ATCC (catalog no. CRL-2500). Wild-type Madin-Darby canine kidney (MDCK) cell line is a kind gift from Dr. Chan Young Park lab, Harvard school of public health. This cell line has been reported in several publication from the group mentioned above. Cells were cultured in low glucose DMEM (Gibco catalog no 11054-001) in presence of 10% FBS (Gibco catalog no. 10437-028) in standard T75 flasks (BD) in a humidified incubator kept at 37°C. For imaging experiments, cells were dislodged from the T75 flask using 0.25% trypsin (Gibco catalog no. 15090-046) with100 μM ETDA solution, and re-plated in 60 mm glass bottom petri plates (Cellvis catalog no. D60-30-1.5-N). For the mitotic cell measurement, cells were allowed to grow 2 days after plating before imaging. Hoechst 33342 (Thermo scientific catalog no. R37605) and CellMask deep red plasma membrane stain (ThermoFisher, cat no. C10046) were added 30 min before imaging. Senescent cells were prepared by plating cells on glass bottom petri-plates as described above. Cells were allowed to adhere to the glass surface overnight and Doxorubicin (100 ng/mL) is added the next morning to induce senescence. Cells were kept under Doxorubicin treatment for 48 Hrs.

#### 3. Animal protocols

Wild type mouse tissues were collected from 8-12 week old male mice in accordance with the procedure approved by Institutional Animal Care and Use Committee (IACUC) at Harvard University. Transgenic female APP-PS1 mice and age-matched wild type (WT) C57 black 6 female mice were purchased from Jackson Laboratory. GDF15-KO mouse tissue was a kind gift from Jui-Hsia Weng. This mouse strain is, without toxin challenge, indistinguishable from WT for our purpose. For live imaging of zebrafish embryos, AB strain wild type fish was used. All fish were kept at 28˚C on a 14-hour-light/10-hour-dark cycle. Embryos were collected from natural crosses. The chorion was removed manually prior to imaging. All fish-related procedures were carried out with the approval of Institutional Animal Care and Use Committee (IACUC) at Harvard University. All animal experiments were approved by the Institutional Animal Use and Care Committee at Harvard Medical School and Massachusetts General Hospital.

#### 4. Tissue sample preparation

All mouse tissues were dissected from euthanized animals and immersion fixed in 4% formaldehyde at 4°C for 24 hours. Mouse cerebellum was perfusion fixed with 4% formaldehyde at 4°C for 24 hours. The fixed tissues, except for cartilage and APP-PS1 and WT brains, were embedded in 2% agarose and sectioned to 40–100 μm thickness using a vibrating microtome (Precisionary Instruments, VF-300-0Z). Fixed cartilage tissues were embedded in BSA-gelatin gel cured with formaldehyde in room temperature overnight, and sectioned by a vibrating microtome (Leica, VT1000M). Fixed brains were transferred into 30% sucrose at 4°C until the tissue sunk in. Then the tissue was frozen in OCT to be sectioned into 25-μm slices in a cryostat. The brain sections were washed of sucrose with phosphate buffered saline. The tissue sections were transferred to phosphate buffered saline bath and stored at 4°C until imaging. For imaging, a tissue section was sealed between a coverglass and a slide glass with phosphate buffered saline using a double-sided tape spacer (Grace Biolabs, SS1X13) or using nail polish as sealant.

#### 5. Tomographic phase microscopy imaging protocol

For tomographic phase imaging, HeLa cells were cultured in a glass bottom dish (Cellvis, D60-30-1.5-N) using a culture insert (Ibidi, 80209) to restrict growth area. The cells were fixed with 4% formaldehyde in the room temperature for 20 minutes and washed with phosphate buffered saline 3 times. A top coverslip is placed over the fixed HeLa cells with an imaging spacer of 120 μm thickness (Grace Biolabs, SS1X20). Tomographic phase images were acquired using a commercial tomographic phase microscope (Tomocube, HT-2) using a 60x water immersion objective lens and a water immersion condenser lens. The dry mass concentration was calculated from the refractive index tomogram assuming average refractive index increment of dn/dc = 0.19 ml/g[9].

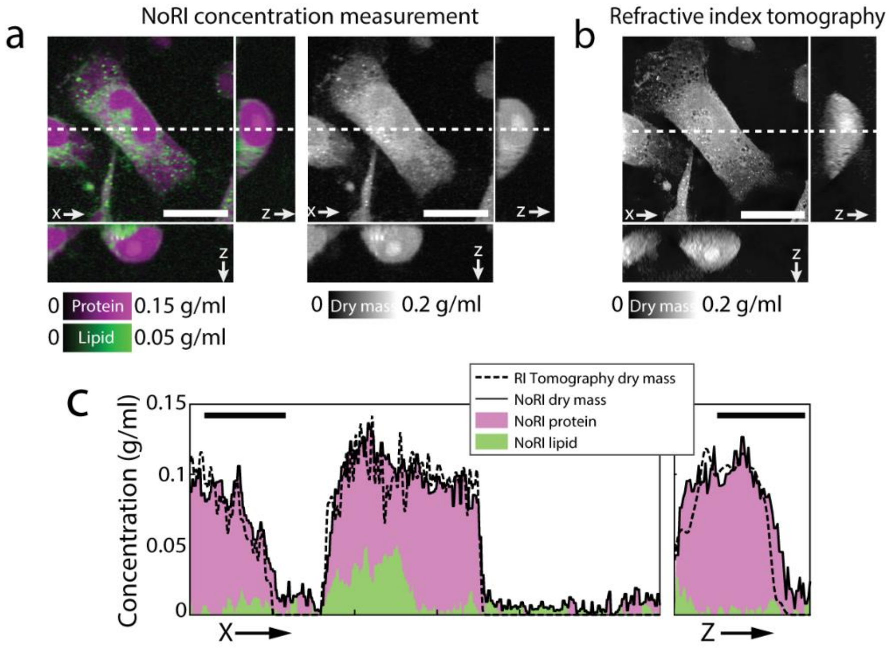


**Supplementary Figure 12.** Comparison of NoRI measurement and Tomographic Phase microscopy (a) Protein and lipid NoRI image of a fixed HeLa cell. Dry mass is the sum of protein and lipid masses. Scale bar, 20 µm. (b) Dry mass concentration of the same cell measured by refractive index tomography microscope (Tomocube, HT-2). Average refractive index increment of 0.19 ml/g is assumed to compute dry mass concentration from refractive index contrast1. Scale bar, 20 µm. (c) Mass concentration profile along the x- and z-cross sections marked by dashed lines in (a) and (b). Dry mass density measured by NoRI is in good agreement with dry mass measured by refractive index tomography. Pearson correlation coefficient of NoRI dry mass density and refractive index tomography dry mass density is r=0.736. Scale bar, 10 µm.

#### 6. Image segmentation method

##### 6.1. Image segmentation based on protein and lipid concentrations

Differences in protein and lipid concentrations were used for segmentation of cell body, nucleus, and lipid droplets in Fig. 2c. To segment the cell body, threshold of 0.0217 g/ml was applied to the protein concentration image, followed by 'Fill Holes' and 'Despeckle' functions of ImageJ. To segment the nucleus, 2-fold scaled lipid image was subtracted from the protein image, from which nuclear area was segmented by thresholding operation. Segmentation was cleaned up by removing areas under 100 $\mu m^2$ in size. Lipid droplets were segmented by threshold operation on lipid concentration. The lipid droplet threshold was adjusted between 0.1-0.2 g/ml depending on the cytosolic lipid concentration (Supplementary Fig. 12). Whether protein and lipid concentration provide enough contrast at the cell body boundary depended on the cell and tissue types. When simple thresholding operation was not sufficient, we found that ImageJ plug-in trainable WEKA segmentation provides a more flexible solution for image segmentation of cultured cells[10]. Single cell segmentation in complex tissue environment can be more challenging and may require optimization of machine learning algorithm or use of fluorescence labels.

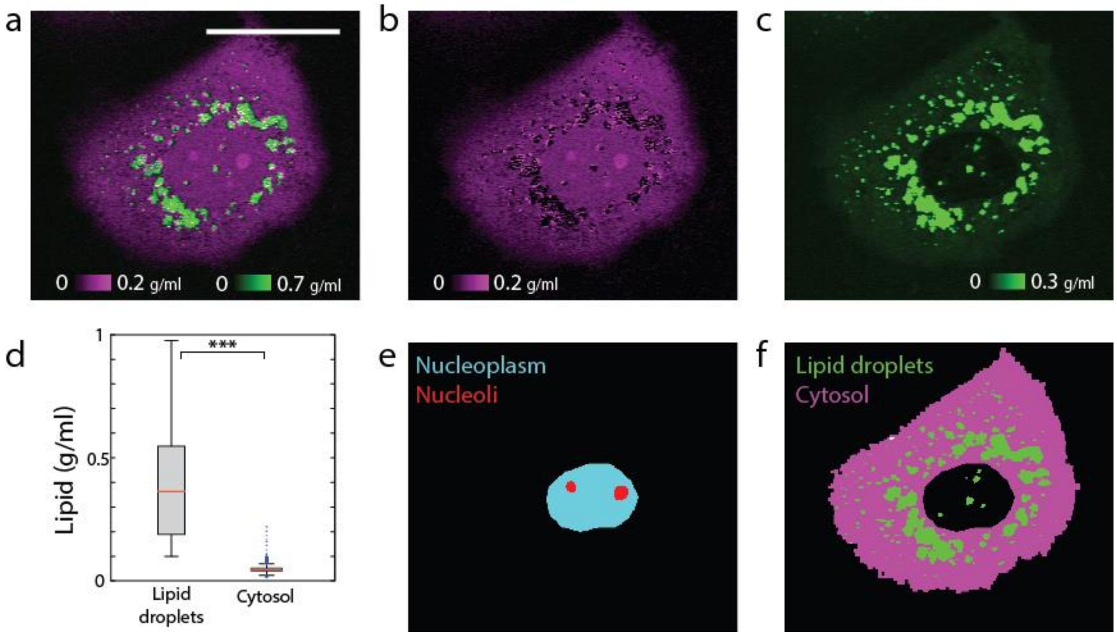


**Supplementary Figure 13. X-Y cross section image and organelle segmentations of a senescent MDCK cell.** (a) Protein concentration (magenta) and lipid concentration (green). Lipid droplets are clearly visible. Scale bar, 50 μm. (b) Protein concentration. (c) Lipid concentration. Color display range for lipid is zoomed in to highlight the low concentration lipid in the cytosol compartment. (d) Distribution of lipid concentrations of lipid droplets (0.387±0.208 g/ml) and cytosol (0.047±0.013 g/ml) in the cell. The large difference facilitates lipid droplet detection ($p<0.001$). (e) Segmentation of nucleolus by trainable WEKA segmentation of nucleus protein image. (f) Lipid droplet and cytosol segmentation by thresholding the lipid image.

##### 6.2. Image segmentation of brain tissue NoRI images

Protein aggregates (amyloid plaques) and lipid aggregates are detected from NoRI images of APP-PS1 brain tissue based on protein and lipid concentration using Trainable WEKA Segmentation tool[10] and OrientationJ[11] in imageJ. Using the WEKA tool, protein concentration image is classified using two classes, protein dense plaques and the rest which includes rest of the tissue and background areas. Lipid concentration image is classified using three classes, white matter which has the highest concentration of lipid, neuropil of the gray matter, and lipid aggregates that show higher lipid concentration compared to the neuropil but lower lipid concentration than white matter. Protein plaques segmentation is completed by applying size threshold of 200 μm$^2$ to the WEKA segmentation output of protein dense plaque class. Lipid aggregates segmentation is calculated from the WEKA segmentation output of lipid aggregate class after removing crosstalk from the myelin class. The crosstalk appears at the border between the two classes owing to the imperfect WEKA classifier training, and can be improved by subtracting myelin adjacent area computed from OrientationJ operation. OrientiationJ detects spatial coherence of features and is particularly suitable to detect the cable-like morphology of myelin sheaths. Myelin-adjacent area is computed by applying threshold operation to spatial coherence output of OrientationJ, and this mask is subtracted from lipid aggregate class segmentation. To obtain the final segmentation for lipid aggregates, size threshold of 35 μm$^2$ is applied. Myelinated axons are segmented by lipid concentration being 0.148 g/ml and higher. Cell body is segmented by lipid concentration threshold at 0.041 g/ml. Neuropil segmentation is computed from the WEKA neuropil class output by subtracting the areas of cell body, amyloid plaques and myelin.

#### 7. Statistics

All statistical analyses were performed in Matlab (R2020a). Box plots represent the median, 25th and 75th percentiles and whiskers with staples represent the minimum and maximum of non-outlier samples. Outliers are plotted as black filled circles. Violin plot is plotted using VIOLIN.m in Matlab[12]. Bars inside violin plots represent the median. Correlations between two distributions and corresponding p-values were computed using CORRCOEF.m function in Matlab. Comparisons between two distributions were performed with two-sample *t*-test using TTEST2.m function in Matlab.

#### 8. Supplementary Theory of NoRI

The linear dependency of SRS intensity to the analyte concentration can be described by an equation $\mathsf{M}_{ij}^{(k)}\mathsf{S}_{ik} = \mathsf{C}_{jk}$(Eq. 1), where $\mathsf{S}_{ik}$ is the SRS signal at the *i*-th Raman band (*i*=$CH_3$, $CH_2$ and $H_2O$) of the *k*-th reference sample (*k*=BSA, DOPC and water samples) and $\mathsf{C}_{jk}$ is the volume concentration of the *j*-th component (*j*=protein, lipid and water) of the *k*-th sample in the unit of volume fraction (ml/ml)[13]. $\mathsf{M}_{ij}^{(k)}$ is the decomposition matrix for the *k*-th sample. In Supplementary Methods 1.5 we described the optimal protocol for measuring the SRS intensity of the calibration reference samples. This minimizes sources of intensity variation other than the sample's chemical composition and the intrinsic Raman cross sections of the chemical components and assures quantitative consistency of the decomposition matrix $\mathsf{M}_{ij}$ across different samples (*k*=BSA, DOPC and water samples). This allows us to concatenate the equations $\mathsf{M}_{ij}\mathsf{S}_{ik} = \mathsf{C}_{jk}$ for all *k* samples. Then, the decomposition matrix is solved by a simple matrix inversion $\mathsf{M}_{ij} = \sum_k \mathsf{C}_{jk}\mathsf{S}_{ik}^{-1}$ (Eq. 2). The inverse of the decomposition matrix $\mathsf{M}_{ij}^{-1}$ is interpreted as the SRS intensity of the *i*-th Raman band measured from a pure material of the *j*-th component. It is proportional to the Raman cross section of the molecule $\sigma_{ij}$ as well as to the efficiency of the imaging system $\mathsf{A}_{i0}$ as optimized for the imaging of calibration standards; that is, $\mathsf{M}_{ij}^{-1} = \sigma_{ij}\mathsf{A}_{i0}$, or $\mathsf{S}_i = \sum_j \sigma_{ij}\mathsf{A}_{i0}\mathsf{C}_j$ (Eq. 3). While the Raman cross section $\sigma_{ij}$ is an intrinsic property of the chemical constituent, the efficiency of the microscope $\mathsf{A}_{i0}$ is affected by the optical aberration of the imaging system and the detector efficiency. Spectral decomposition is applied by matrix multiplication of the decomposition matrix with SRS images at the $CH_3$, $CH_2$ and $H_2O$ Raman bands: $\sum_i \mathsf{M}_{ij}\mathsf{S}_i(\vec{r})$ (Eq. 4).

When imaging samples, the light scattering and aberration caused by the sample introduces an extra attenuation factor $\mathsf{A}_X(\vec{r})$ to the signal, which can be expressed as $\mathsf{S}_i(\vec{x}) = \sum_j \sigma_{ij}\mathsf{A}_{i0}\mathsf{A}_X(\vec{r})\mathsf{C}_j(\vec{r})$ (Eq. 5). As we argue below, we assume that the sample induced attenuation factor is independent of the Raman bands. When spectral decomposition is applied to sample images, by multiplying the decomposition matrix with the SRS intensity, we do not get the true concentration because of this factor. Instead, the output of spectral decomposition is proportional to the concentration of respective chemical components and also to the spatially heterogeneous attenuation: $\sum_i \mathsf{M}_{ij}\mathsf{S}_i(\vec{r}) = \mathsf{A}_X(\vec{r})\mathsf{C}_j(\vec{r})$ (Eq. 6). The unknown attenuation factor varies with position and with each sample and limits the quantitative interpretation of spectral decomposition. The sample attenuation $\mathsf{A}_X(\vec{r})$ can be estimated by making a reasonable approximation that the major chemical components (protein, lipid and water) occupy 100% of the sample volume. This is expressed by $\sum_j \mathsf{C}_j = 1$ (Eq. 7) where *j*=protein, lipid and water. Using Eq. 7, the sum of Eq. 6 over *j*=protein, lipid and water is equal to $\mathsf{A}_X(\vec{r}) = \sum_{ij} \mathsf{M}_{ij}\mathsf{S}_i(\vec{r})$ (Eq. 8). $\mathsf{A}_X(\vec{r})$ measures the collective sample property that attenuates the SRS intensity at each voxel which includes the presence of scatterers above and below the imaging plane (Supplementary Fig.7f). It is defined throughout the sample imaging volume and serves as the light scattering normalization mask. Absolute concentration of *j*-th component in volume fraction $\mathsf{C}_j(\vec{r})$ is obtained by dividing the spectral decomposition in Eq. 6 by the normalization mask $\mathsf{A}_X(\vec{r})$ from Eq. 8. Due to the general nature of the linear relationship in Eq. 5, light scattering normalization may be applied to other nearby Raman bands provided that the light scattering property of the laser does not change significantly between the applied band and the $CH_3$, $CH_2$ and $H_2O$ bands. Additional information on the wavelength dependence of tissue light scattering in the near-infrared range is included in the following section.

#### 9. Wavelength dependence of light scattering: impact on imaging depth limit

NoRI algorithm corrects for the effect of light scattering on the basis that the light scattering is same for all pump beam wavelengths. In fact, the intensity attenuation from tissue light scattering and absorption is nearly independent of wavelength in the relevant wavelength range[14]. Pump beam wavelength changes between 770 nm and 805 nm to select different Raman bands. The intensity $I$ at depth $z$ in a tissue with reduced scattering coefficient $\mu'_s(\lambda)$ is described by $I(z,\lambda) = I_0 e^{-\mu'_s(\lambda)z}$. Reduced scattering coefficient of typical tissue in 770–805 nm is on the order of 1-2 mm$^{-1}$, and wavelength dependent change of this value is less than 0.2 mm$^{-1}$ between 770–805 nm[15]. The wavelength dependent difference in light scattering can be described using $I(z,\lambda_1)/I(z,\lambda_2) = I_0 e^{-(\mu'_s(\lambda_1)-\mu'_s(\lambda_2))z}$. The depth $z$ where the difference reaches 1/e is $z = 1/(\mu'_s(\lambda_1) - \mu'_s(\lambda_2))$. Similarly, the depth $z$ where the attenuation deviates by 1% is $z = \log(1.01)/(\mu'_s(\lambda_1) - \mu'_s(\lambda_2))$. We simulated the imaging depth at which the wavelength dependent differences in reduced scattering length affect 1% and 2% of intensity using the average number reported in Ref[15] (Supplementary Table 2). Various tissues show less than 2% difference between 770 and 805 nm within the imaging depth limit of SRS microscopy (90-120 μm)[16].

**Supplementary Table 2. Wavelength dependence of light scattering of various tissues**

| Tissues | $\mu'_s(\lambda)$ (mm$^{-1}$) | | | | $\mu'_s$(770nm) $-\mu'_s(805nm)$ (mm$^{-1}$) | 1/e difference depth (mm) | 1% difference depth (mm) | 2% difference depth (mm) |
|---|---|---|---|---|---|---|---|---|
| | $\lambda$=500 nm | $\lambda$=770 nm | $\lambda$=800 nm | $\lambda$=805 nm | | | | |
| Skin | 4.6 | 2.49 | 2.36 | 2.34 | 0.15 | 6.6 | 0.028 | 0.056 |
| Brain | 2.42 | 1.21 | 1.13 | 1.12 | 0.08 | 12.0 | 0.052 | 0.103 |
| Breast | 1.68 | 1.07 | 1.02 | 1.02 | 0.05 | 20.5 | 0.089 | 0.176 |
| Bone | 2.29 | 1.68 | 1.64 | 1.63 | 0.05 | 19.0 | 0.082 | 0.163 |
| Other soft tissues | 1.89 | 1.08 | 1.03 | 1.02 | 0.06 | 16.6 | 0.072 | 0.143 |
| Other fibrous tissues | 2.71 | 1.34 | 1.26 | 1.25 | 0.09 | 10.7 | 0.046 | 0.092 |
| Fatty tissue | 1.84 | 1.38 | 1.34 | 1.34 | 0.04 | 24.7 | 0.107 | 0.212 |

#### 10. Depth dependence of concentration measurement sensitivity

Here we simulated the effect of signal to noise ratio (SNR) on the sensitivity of concentration calculation. Especially, SNR decreases as a function of imaging depth as is depicted in Supplementary Fig. 13a. The detector noise was 26 nA as measured by the standard deviation of photo detector readout when the detector is illuminated with pump beam and Stokes beam is absent from the sample. SRS signal of tissue samples is around 2 μA (SNR=100) near the surface and exponentially decreases with imaging depth. 1/e$^2$ attenuation depth of various murine tissues are in the range of 90 - 120 μm[16]. We used 100 μm for our simulation. The noise was generated from a normal distribution with 0 nA mean and standard deviation of 26 nA. As the noise in the $CH_2$, $CH_3$ and $H_2O$ Raman bands are independent, we denote the three instances of noise by a 3x1 vector $\vec{n}$. Spectral decomposition transforms the detector signal $\vec{s} + \vec{n}$ to $M(\vec{s} + \vec{n})$, where $\vec{s}$ is true signal in the $CH_2$, $CH_3$ and $H_2O$ Raman bands and $M$ denotes an experimentally measured decomposition matrix. $M\vec{s}$ is a 3x1 vector whose components are proportional to the true concentration in protein, lipid and water fractions ($\vec{c}$). There is an arbitrary unit constant ρ that satisfies $M\vec{s} = \rho\vec{c}$ for all $\vec{s}$. $M\vec{n}$ is a 3x1 vector whose components represent the noise in protein, lipid and water fractions. They had mean value of 0 and standard deviations of 20, 26, and 28 (A.U.) for protein, lipid, and water fractions, respectively. The normalization step transforms $M\vec{s} + M\vec{n}$ to absolute concentration in volume fraction $\vec{C} = \frac{M\vec{s}+M\vec{n}}{\sum(M\vec{s}+M\vec{n})}$, where the sum in the denominator is over protein, lipid and water fractions. Note that when SNR equals 1, the mean of $Ms_i$ equals the RMS size of $\langle Mn_i \rangle$, for each $i$=protein, lipid or water. Therefore, we can obtain $\rho c_i = \langle Mn_i \rangle$. Since $c_i$, the concentration in volume fraction, is in the range of 0 - 1, we can simulate different SNR by parameterizing $\rho$. We computed the mean and RMS noise of the absolute concentration $\vec{C}$ for concentration range of 0 – 1 (v/v) and SNR range of 1 to 100 at steps of 5 (Supplementary Fig. 13b-c). Volume fraction is converted to w/v concentration using the density of water (1 g/ml). The sensitivity at 100 SNR was 0.015 g/ml (1.5% w/v) in agreement with the experimental measurement. At the 1/e attenuation depth (50 μm in this simulation), the noise reaches 0.040 g/ml (4% w/v) for single pixels and 0.020 g/ml (2% w/v) for average of 4 pixels. By averaging 64 pixels (equivalent to 3.3 μm x 3.3 μm area), noise can be kept under 0.015 g/ml (1.5% w/v) at the 1/e$^2$ attenuation depth (100 μm).

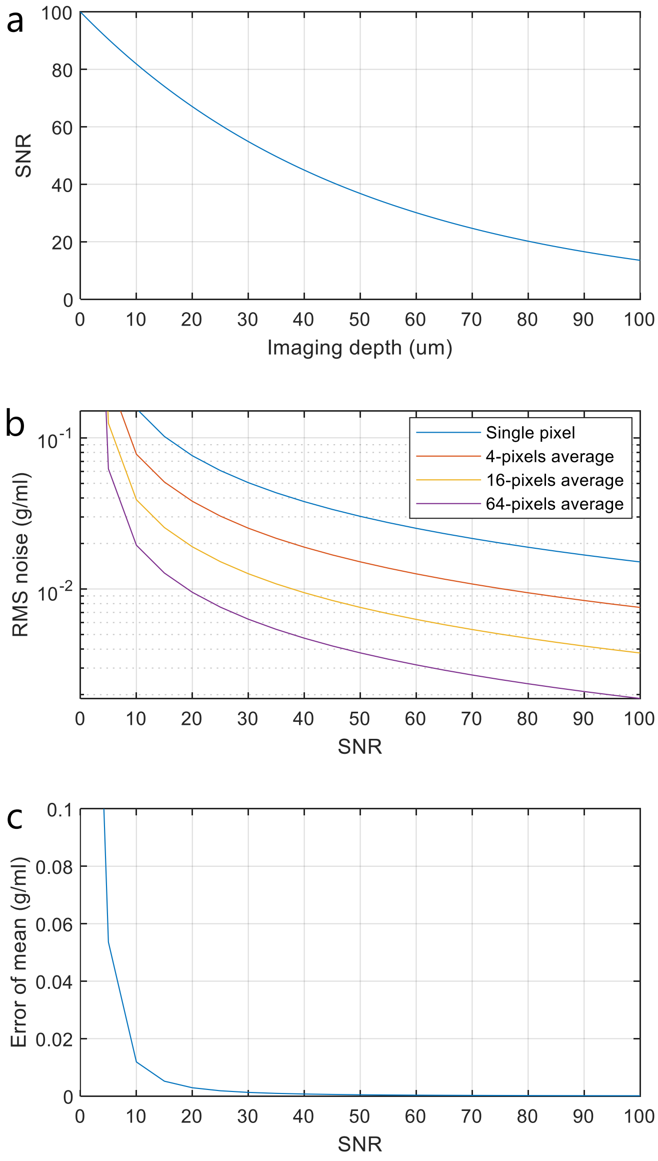


**Supplementary Figure 14. Sensitivity of concentration as a function of signal-to-noise ratio (SNR).** (a) SNR decreases with imaging depth. (b) RMS noise as a function of SNR. (c) Error of mean value as a function of SNR.

#### 11. Limitation of the absolute concentration measurement by NoRI

To assure the accuracy of the absolute concentration measurement by NoRI, its application should be applied to samples where the following conditions are satisfied: (11.1) protein, lipid and water constitute most of the chemical composition, (11.2) calibration standards (BSA and DOPC) are good approximations of the average proteome and lipidome of the sample in terms of their Raman spectra, and (11.3) pigments are absent or bleached as they can cause non-Raman background through two-color two-photon absorption of pump and Stokes beams[17,18].

##### 11.1. Chemical components of sample

NoRI based on the $CH_3$, $CH_2$ and $H_2O$ channels provides high accuracy for cells and tissues where protein, lipid and water makes up nearly 100% of the chemical composition. For mammalian cells whose combined fraction of protein, lipid and water is greater than 93% of the wet mass[19], the percentage error arising from neglected 7% other component is small in absolute size for components other than water because it is multiplicative (7% error of 0.1 g/ml is ± 0.007 g/ml). For tissues with very low lipid content, NoRI normalization scheme can be adapted to use only the $CH_3$ and $H_2O$ channels. Such approach was used to analyze water content in eye lens tissue by confocal Raman microscopy [20]. Normalization by NoRI scheme can be generalized to more than 4 chemical components to improve accuracy when the sample has significant amount of other substances (Supplementary Fig. 15). For example, yeast and bacterial cells have high nucleic acids content[19], and 4-channel NoRI including nucleic acids will be more appropriate. Increasing the number of Raman bands will also increase acquisition time and will benefit from hyperspectral SRS microscopy and advanced spectral separation schemes[21,22].

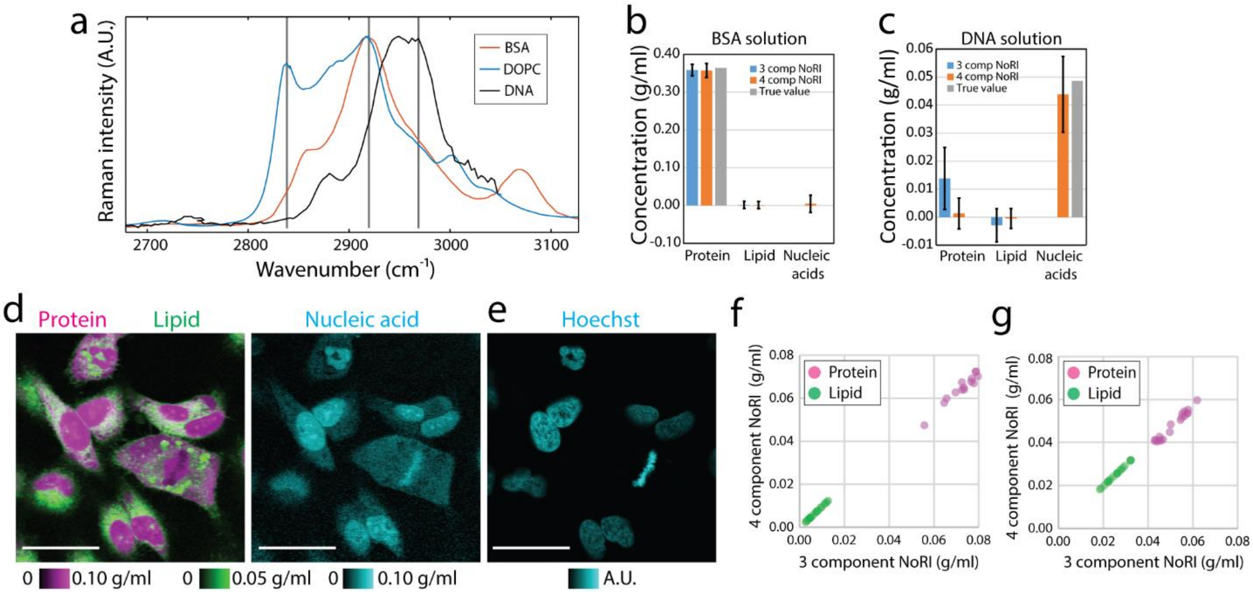


**Supplementary Figure 15. 4-component NoRI for quantitative measurement of protein, lipid, nucleic acids, and water.** (a) Raman spectrum of BSA, DOPC and salmon sperm DNA. The contribution from solvents is subtracted and the effect of pulse overlap is normalized according to Supplementary Fig. 3. Vertical lines indicate the positions of the $CH_2$, $CH_3$, and nucleic acids bands from the left to the right. These three Raman bands are measured along with the $H_2O$ band to spectrally separate proteins, lipids, nucleic acids and water. The laser setting for selecting these bands are summarized in Supplementary Table 1. (b-c) Comparison of concentration measurement of pure samples by 3-component NoRI vs 4-component NoRI. DNA shows partial cross talk with protein measurement. (d) NoRI imaging of protein, lipid, and nucleic acids in live HeLa cells measured by 4-component NoRI. Raw SRS images are acquired using frame averaging of 16 to suppress detector noise. Scale bar, 40 µm. (e) Sample is co-stained with Hoechst 33342 and imaged using confocal scanning microscopy. Nucleic acid rich area in the NoRI image co-localizes with nuclei in the Hoechst fluorescence image. (f-g) Comparison of protein and lipid measurement in HeLa cells by 3-component NoRI vs 4-component NoRI in nucleus (f) and in cytoplasm outside nucleus (g) (N=18).

##### 11.2. Calibration reference materials

We chose BSA and DOPC as the calibration reference materials because BSA has a similar Raman spectrum in the $CH_2$ and $CH_3$ bands compared to the average vertebrate proteome[23] and DOPC is one of the most abundant lipid in the cell membranes. They are economical and available as a highly purified material in large quantity, and as discussed below, are generally good starting material when considering the choice of calibration standards.

Whether BSA is a good representation of sample proteome is solely determined by their high wavenumber Raman spectrum. This can be estimated from the hydrocarbon frequency of the sample proteome which can be counted from the amino acid frequency. As shown in Supplementary Fig. 16, the frequency of methyl groups, methylene groups and other hydrocarbons of BSA is similar to that of the average vertebrate proteome[24]. When a few types of proteins dominate the tissue proteome, it should be checked whether BSA is a good representation of sample proteome's Raman spectrum. In such cases, the dominant protein or tissue protein extract can be used as the calibration material[23].

**Supplementary Table 3. Frequency of methyl and methylene groups in the average vertebrate proteome[24] and in bovine serum albumin.** BSA amino acid frequency from https://www.uniprot.org/uniprot/P02769

| Amino acids | Amino acids | $CH_3$ in side group | $CH_2$ in side group | Other C-H bonds | Observed Frequency in Vertebrates | Frequency in BSA |
|---|---|---|---|---|---|---|
| Alanine | A | 1 | | | 7.4% | 8.1% |
| Arginine | R | | 3 | | 4.2% | 3.9% |
| Asparagine | N | | 1 | | 4.4% | 2.4% |
| Aspartic Acid | D | | 1 | | 5.9% | 6.9% |
| Cysteine | C | | 1 | | 3.3% | 6.0% |
| Glutamic Acid | E | | 2 | | 5.8% | 10.1% |
| Glutamine | Q | | 2 | | 3.7% | 3.4% |
| Glycine | G | | | 1 | 7.4% | 2.7% |
| Histidine | H | | 1 | 2 | 2.9% | 2.9% |
| Isoleucine | I | 2 | 1 | 1 | 3.8% | 2.4% |
| Leucine | L | 2 | 1 | 1 | 7.6% | 10.5% |
| Lysine | K | | 4 | | 7.2% | 10.1% |
| Methionine | M | 1 | 2 | | 1.8% | 0.7% |
| Phenylalanine | F | | 1 | 5 | 4.0% | 4.6% |
| Proline | P | | 2 | | 5.0% | 4.8% |
| Serine | S | | 1 | | 8.1% | 4.8% |
| Threonine | T | 1 | | 1 | 6.2% | 5.7% |
| Tryptophan | W | | 1 | 5 | 1.3% | 0.3% |
| Tyrosine | Y | | 1 | 4 | 3.3% | 3.4% |
| Valine | V | 2 | | 1 | 6.8% | 6.2% |

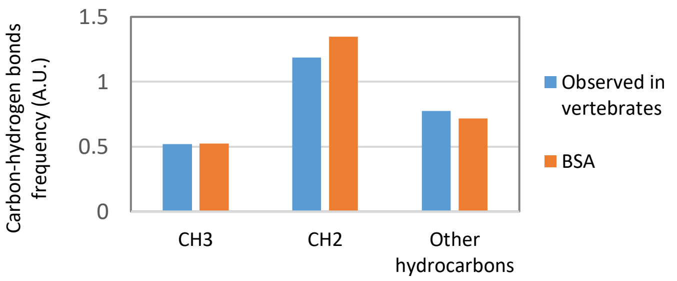


**Supplementary Figure 16. Carbon-hydrogen bonds frequency in vertebrate proteome and bovine serum albumin.** Bonds frequency are calculated from Supplementary Table 3.

Although DOPC is the most abundant lipid in cell membranes, lipid droplets are composed of triacylglycerol and cholesterol esters, and spectral decomposition using DOPC reference spectra does not calculate the accurate lipid concentrations in the lipid droplets. As discussed above, we adopted a two-step approach in which lipid droplets are segmented by thresholding operation on lipid concentration (Supplementary Fig. 13). The lipid in the aquatic phase of cytoplasm (lipids that are not in lipid droplets) would be composed of membrane lipids and are better represented by DOPC. Alternatively, one can perform multi-component NoRI using appropriate Raman bands to quantify different types of lipids as separate components, as saturated lipids, unsaturated lipids, cholesterol and proteins in biological samples can be spectrally unmixed by using 2800-3100 $cm^{-1}$ Raman bands[2,22,25–27]. Even for the membrane lipids, enrichment of different types of lipids may cause the sample lipidome spectrum deviate from the reference spectrum of DOPC[2,28,29]. In such cases, one can achieve better spectral decomposition by using tissue lipid extract as the calibration standard.

##### 11.3. Pigmented samples

SRS overcame the large part of non-Raman background of CARS[1,30], and non-Raman backgrounds from cross-phase modulation (XPM) and thermal lensing can be negligible in comparison with the strong Raman signals from the $CH_2$, $CH_3$ and $H_2O$ bands (Fig. 1c)[31–33]. The ratio of signal to background can decrease with depth in thick or highly scattering samples, when the sample configuration requires low NA collection optics, or when imaging weak Raman bands in the fingerprint region or low concentration Raman probes. In such applications, it is a standard approach to subtract the non-Raman signal from Raman peak images to increase accuracy[34], which can also be employed in NoRI. Since the images in the $CH_3$, $CH_2$ and $H_2O$ Raman bands are much brighter than the non-Raman background, we decided that it is unnecessary to subtract the non-resonant image when measuring unpigmented samples (Supplementary Fig. 17).

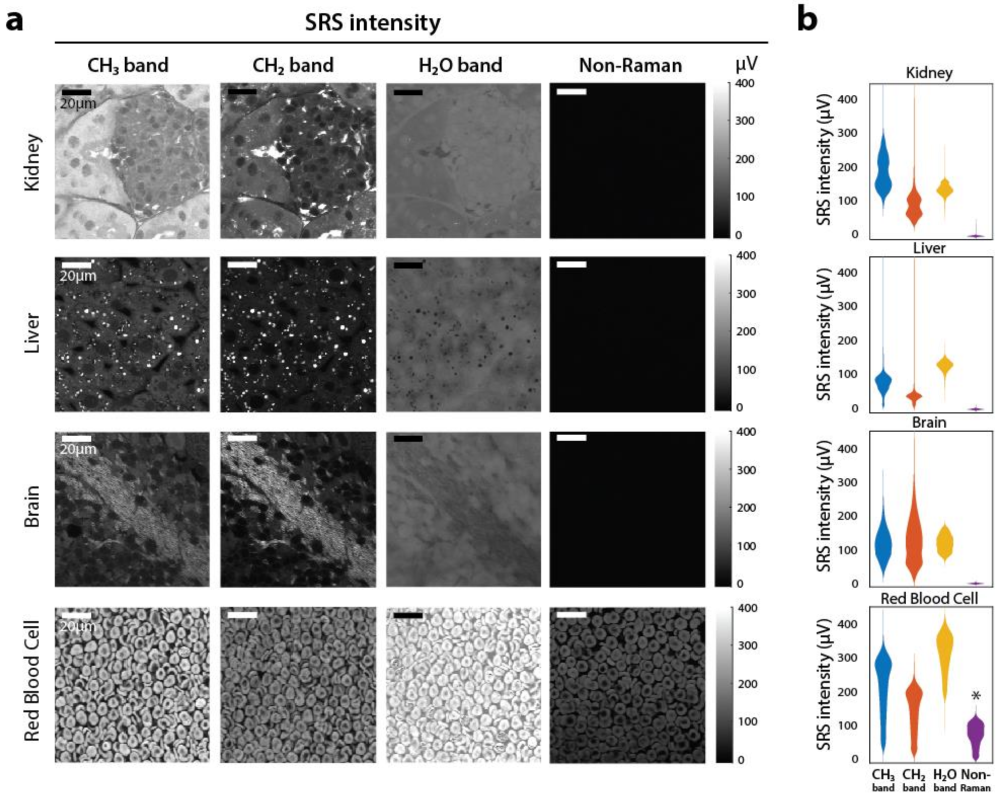


**Supplementary Figure 17. Non-Raman background in various tissue samples and blood.** (a) Non-Raman signal is negligible compared to the SRS signal in the $CH_3$, $CH_2$ and $H_2O$ bands in fixed mouse kidney, liver and brain, but it is large in red blood cells. (b) Quantification of SRS intensity and non-Raman signal intensity from images in (a). Photo current is converted to voltage using a 50 Ω resister.

On the other hand, pigments such as hemoglobin, melanin, and lipofusin can generate two-photo absorption (TPA). Unlike XPM and thermal lensing effect, non-Raman signal from TPA can generate large signal (red blood cells in Supplementary Fig. 17) but it can be subtracted as well due to its flat spectrum[33]. In practice, we removed red blood cells from samples by perfusing the tissue with saline, and bleached melanin with depigmenting buffer (hydrogen peroxide and formamide) as needed to eliminate TPA instead of computationally subtracting them. Even when saline perfusion is imperfect, presence of red blood cells does not interfere with the NoRI measurement of the surrounding tissue. Liver may contain bile pigments such as bilirubin and biliverdin. However, we find that a healthy murine liver tissue from 3 month old mouse showed low level TPA under our imaging condition using wavelengths of 770-805 nm and 1045 nm. This is consistent with the absorption spectrum of bile that rapidly drops over 700 nm[35].

#### 12. Supplementary Figures

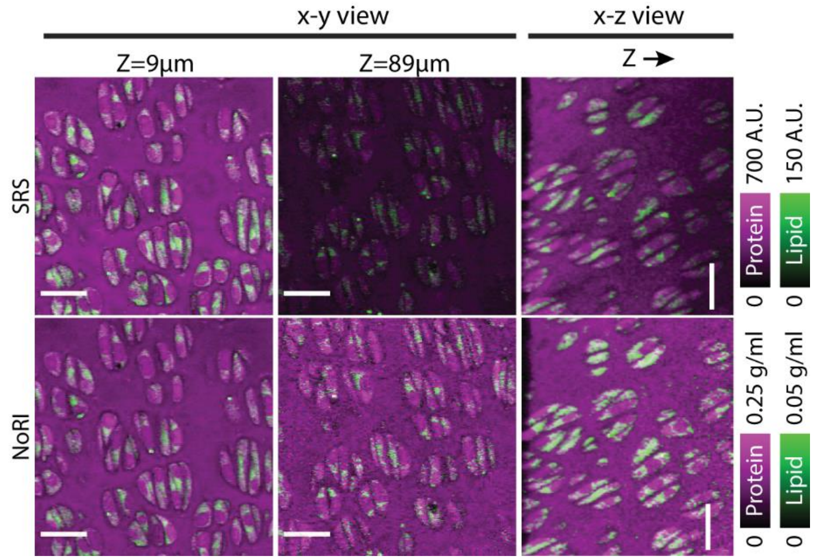


**Supplementary Figure 18.** Protein and lipid distribution image of live mouse cartilage tissue before and after NoRI normalization. Signal intensity diminishes with imaging depth. Scale bar, 20 μm. NoRI normalization removes the effect of light scattering and enables accurate measurement of protein and lipid concentration throughout tissue volume.

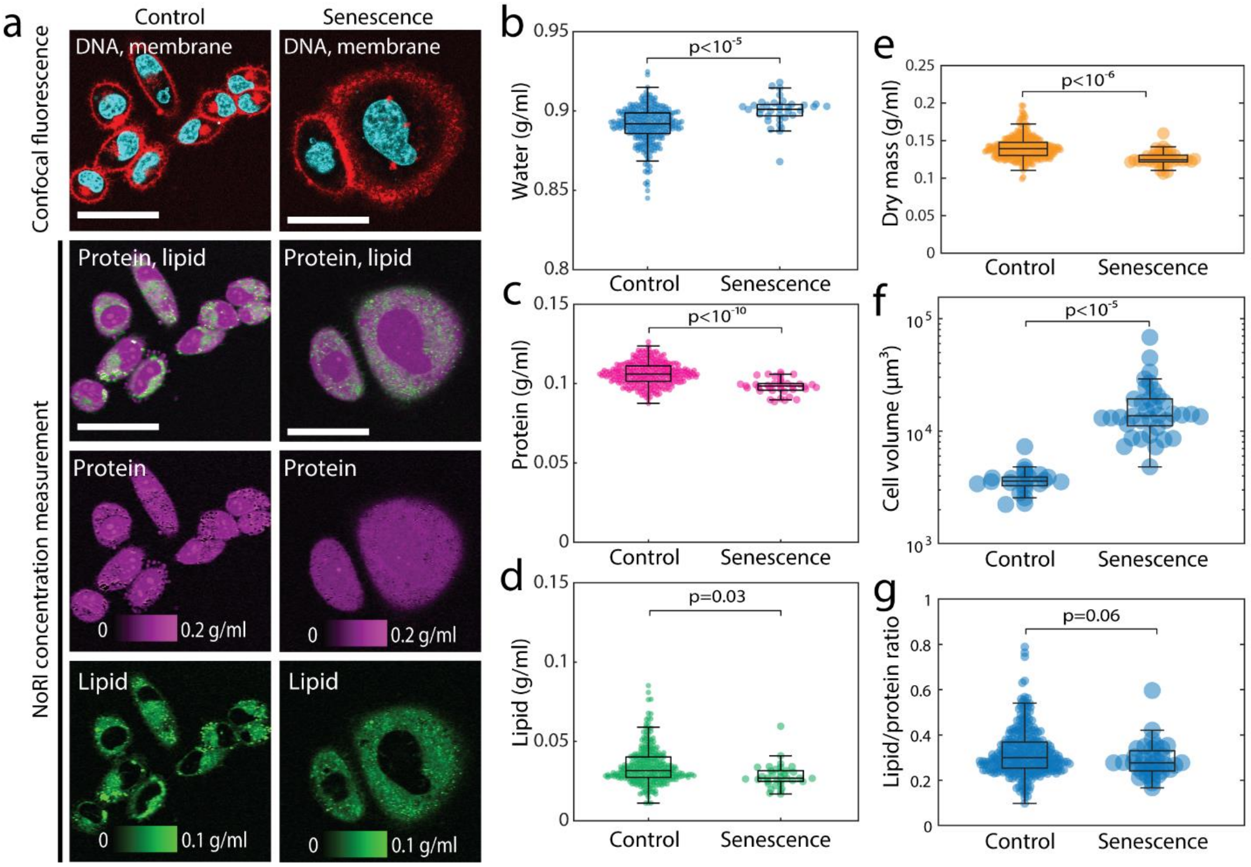


**Supplementary Figure 19. Cytoplasm dilution and hypertrophy of senescent A7 cells.** (a) Representative images of proliferating and senescent live A7 cells. Senescence is induced by 48 hour doxorubicin treatment. Scale bar, 50 μm. (b-e) Senescent A7 cells have lower protein, lipid, and dry mass concentration and higher water concentration compared to proliferating cells. (Control N=286, mean and standard deviation are water 0.891±0.011 g/ml, protein 0.106±0.007 g/ml, lipid 0.035±0.012 g/ml, dry mass 0.141±0.015 g/ml. Senescence N=35, mean and standard deviation are water 0.900±0.009 g/ml, protein 0.098±0.005 g/ml, lipid 0.028±0.008 g/ml, dry mass 0.127±0.010 g/ml.) (f) Cell volume increase in senescent cells. Mean cell volume is 3693±1035 fl in control cells and 17289±11893 fl in senescent cells. (p<0.001) Senescent cell volume is measured from membrane localized fluorescence marker (N=36) and control cell volume is measured from 3D segmentation of NoRI dry mass concentration images (N=22). (g) Ratio of lipid mass to protein mass in single cells is 0.32±0.10 in control A7 cells and 0.29±0.08 in senescent A7 cells. (p=0.065)

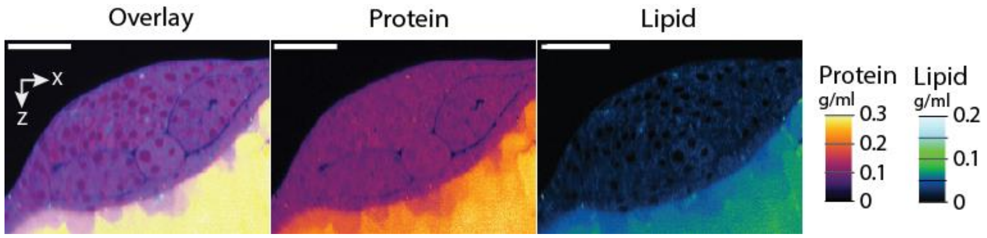


**Supplementary Figure 20. NoRI image of live zebrafish embryo at the 6-somite stage.** Scale bar, 50 μm. The protein and lipid concentrations at this stage did not vary much with different embryonic cell types. As expected, yolk granules are packed with proteins and lipids. Individual yolk granules could be recognized adjacent to the embryo as they had lower protein and lipid concentrations, their stores presumably having been partially depleted in the developmental process.

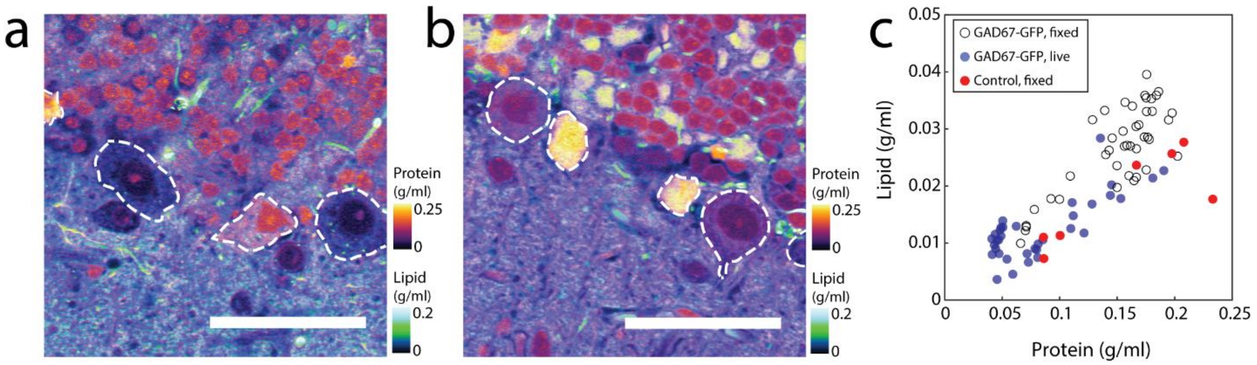


**Supplementary Figure 21** (a) A representative NoRI image of live Purkinje cells (outlined) in brain slice from a GAD67-GFP expressing mouse. Scale bar 50 μm. (b) A representative NoRI image of fixed Purkinje cells (outlined) from a mouse that does not express GAD67-GFP (GDF15-KO strain). Scale bar 50 μm. (c) Protein and lipid concentration of Purkinje cells. Concentration variation is reproduced in both live and fixed cells and in GFP expressing and non-expressing cells. Fixed GFP expressing, N=38. Live GFP expressing, N=35. Fixed non-expressing control, N=7.

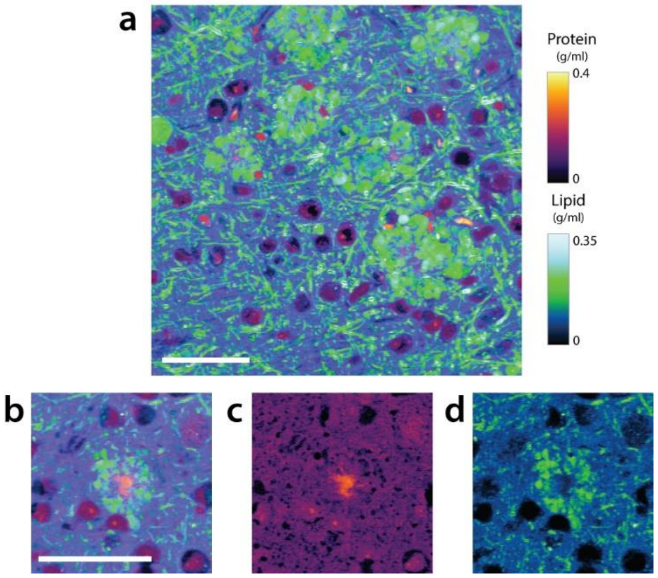


**Supplementary Figure 22. NoRI protein and lipid concentration images of 5xFAD mouse brain tissue.** (a) Protein and lipid concentration images of a 5xFAD mouse brain tissue. (b) Detailed view of a senile plaque in 5xFAD mouse shows a core of protein aggregates surrounded by a lipid-rich corona. Scale bars, 50 μm. (c) Protein only view (b). (d) Lipid only view of (b).
